# Bimodal masked language modeling for bulk RNA-seq and DNA methylation representation learning

**DOI:** 10.1101/2025.06.25.661237

**Authors:** Maxence Gélard, Hakim Benkirane, Thomas Pierrot, Guillaume Richard, Paul-Henry Cournède

**Author notes:** Correspondence to: Maxence Gélard < >.

## Abstract

Oncologists are increasingly relying on multiple modalities to model the complexity of diseases. Within this landscape, transcriptomic and epigenetic data have proven to be particularly instrumental and play an increasingly vital role in clinical applications. However, their integration into multimodal models remains a challenge, especially considering their high dimensionality. In this work, we present a novel bimodal model that jointly learns representations of bulk RNA-seq and DNA methylation leveraging self-supervision from masked language modeling. We implement an architecture that reduces the memory footprint usually attributed to purely transformer-based models when dealing with long sequences. We demonstrate that the obtained bimodal embeddings can be used to fine-tune cancer-type classification and survival models that achieve state-of-the-art performance compared to unimodal models. Furthermore, we introduce a robust learning framework that maintains downstream task performance despite missing modalities, enhancing the model’s applicability in real-world clinical settings.

## 1. Introduction

The growing availability of high-throughput technologies has revolutionized molecular research, generating extensive genomic, transcriptomic, and epigenomic data that hold immense potential for personalized medicine (Ho et al., 2021; Stark et al., 2019; Dai & Shen, 2022). Cancer diagnosis and prognosis thus increasingly rely on heterogeneous patient data, and the integration of these diverse data sources remains a significant challenge, even more so when some modalities may be missing in real clinical applications. The high dimensionality of each modality often makes classic machine learning and deep learning methods ineffective for diagnostic purposes. As a result, there is a growing trend to first learn representations of the data, particularly using self-supervised approaches. In this context, foundation models have steadily emerged as powerful tools to learn effective and generalizable embeddings that can be applied to biological and clinical tasks. Trained with an unsupervised language modeling objective, they have already been applied to a wide range of omics data, including genomics (Dalla-Torre et al., 2025; Brixi et al., 2025), single-cell transcriptomics (Cui et al., 2024) or bulk RNA-seq (Gélard et al., 2025). These models extensively leverage the transformer architecture (Vaswani et al., 2017) which limits the maximum input length of the model due to the quadratic memory scaling of the attention mechanism. Recent models have developed new architectures to cope with these long-range sequences, either by integrating convolutional blocks (Avsec et al., 2021; Linder et al., 2025; Joshi et al., 2025) or state-space models (Popov et al., 2025). Among multiple studies, the Cancer Genome Atlas (TCGA, https://portal.gdc.cancer.gov/) is a publicly available dataset that gathers multi-omics data, in particular bulk RNA-seq and DNA methylation, which are the focus of this work. Including clinical information such as survival time and divided into 33 cohorts (or cancer types), this dataset is a popular benchmark for evaluating survival analysis and cancer-type classification methods. Survival analysis, or time-to-event prediction, aims to predict the time from diagnosis to the patient’s death from the disease using censored data. Though classically tackled with Cox regression (Cox, 1972), the Cox partial likelihood has more recently been reformulated as a loss used to train deep learning architectures (Ching et al., 2018; Katzman et al., 2018).

In this paper, we introduce *MOJO*, standing for **<u>M</u>**ulti-**<u>O</u>**mics **<u>JO</u>**int representation learning, which we tailor in this study for learning joint embeddings of bulk RNA-seq and DNA methylation through bimodal masked language modeling from the TCGA dataset. We leverage a multi-modal architecture that employs a mix of convolutional and transformer layers. We show that the embeddings learned by *MOJO* lead to state-of-the-art performance in various tasks from pan-cancer cancer-type classification, survival analysis, to cancer subtype clustering. We further present a framework that allows for a downstream task model to preserve its performance in the absence of a modality by introducing an auxiliary loss based on mutual information. Additionally, we demonstrate that MOJO’s gene-level aggregation makes it inherently array-agnostic, enabling robust zero-shot transfer to legacy Illumina 27k methylation platforms without any retraining. We also validate out-of-distribution generalization on two fully held-out external cohorts, confirming that its bimodal advantage persists on unseen populations. The code and model weights are publicly available at https://github.com/instadeepai/multiomics-open-research, with pretrained checkpoints on *Hugging Face*.

## 3. Related works

### Omics representation learning

Omics representations have traditionally been derived from statistical methods such as Principal Component Analysis (Jolliffe, 2002) or Non-negative Matrix Factorization (NMF) (Lee & Seung, 1999), the latter being particularly suited for RNA-seq and DNA methylation due to their positivity. Deep learning architectures such as Masked Auto-Encoders (Gross et al., 2024) or Mixture-of-Experts (Meng et al., 2023) have since been applied to learn omics representations used either for survival analysis or cancer-type predictions. In line with foundation models for single-cell transcriptomics (Cui et al., 2024; Yang et al., 2022), Gélard et al. (2025) developed a transformer-based model for bulk RNA-seq representation learning. Multi-modal integration is often performed using late integration; *i*.*e*., each source is encoded separately before being aggregated, often using Kronecker product (Chen et al., 2020), element-wise operations (Vale-Silva & Rohr, 2021) or cross-attention (Garau-Luis et al., 2024). On the other hand, Benkirane et al. (2023) developed a custom integration of multi-omics data using variational auto-encoders (Kingma & Welling, 2013).

### Missing modalities

Improving the robustness of multi-modal models under modality absence is crucial given the sensitivity of recent deep learning architectures to missing modalities (Ma et al., 2022). Part of the literature focuses on techniques that operate at the data level, namely relying on modality imputation (Chen et al., 2024; Zhang et al., 2021a; Ma et al., 2021). Zhi et al. (2024) propose a retrieval-augmented in-context learning framework to address the missing modalities issue in a low-data regime. Another path towards handling missing modalities lies in adjusting the model itself with, for example, model fusion (Wagner et al., 2011) or knowledge distillation (Saha et al., 2024). Training methods are also adapted to make multimodal models robust to missing modalities by employing modality dropout (Krishna et al., 2024; Nezakati et al., 2024) during training to simulate scenarios where a subset of modalities might be missing. In our work, we adapt a technique from Ramazanova et al. (2025), which addresses the challenge of missing modalities as a test-time adaptation problem, by incorporating an auxiliary loss during the fine-tuning of our model.

## 4. Multi-omics joint representation learning

### 4.1. Bulk RNA-seq and DNA methylation processing for language modeling

#### Modalities alignment

Bulk RNA-seq provides an estimate of the mean expression over all cells in a sample for a large number of genes denoted *N*_*genes*_ (typically *N*_*genes*_ ~ 10^4^). Thus, each sample of RNA-seq is composed of real values (in units of transcript per million, or TPM) per gene, 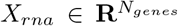, to which we apply a *x* ↣ *log*_10_(1 + *x*) transformation. DNA methylation is the enzymatic attachment of methyl groups to DNA nucleotide bases (usually Cytosine followed by Guanine or *CpG* site). The methylation level of a given site is expressed by a beta value *β* ∈ [0, 1], with the number of measured sites, *N*_*sites*_, being around 450,000, resulting in a methylation sample 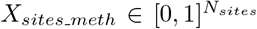. The first step in our modeling is to align RNA-seq and methylation data by defining a methylation value per gene 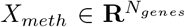. More precisely, for each gene *g* ∈ ⟦1; *N*_*genes*_⟧, we define *sites*(*g*) as all methylation sites associated with the gene, as defined in the Infinium Human Methylation 450k (Bibikova et al., 2011) BeadChip annotations. We evaluate two aggregation strategies for computing gene-level methylation values: (1) a region-weighted strategy where each site *s* ∈ *sites*(*g*) is assigned weight *w*(*s*) based on its genomic region, with higher weights for regulatory regions (Deaton & Bird, 2011), and (2) a simple average where all weights are equal (*w*(*s*) = 1). Both compute gene-level methylation 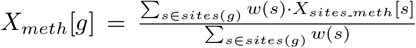. Benchmark experiments in the Appendix demonstrate that the simple average already achieves state-of-the-art performance; thus, we use this method by default. Detailed weight specifications and a comprehensive comparison are provided in Appendix D. Finally, a bimodal sample is represented as 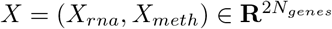.

#### Tokenization

Language models learn to estimate the likelihood of token sequences. Thus, after aligning the two modalities and obtaining a feature vector *X* = (*X*_*rna*_, *X*_*meth*_), each of its components is tokenized by binning their values on linear scales. The token id associated with a given RNA-seq or methylation value is its corresponding bin id. Therefore, after tokenization, one sample is a vector 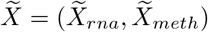 with 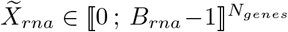 and 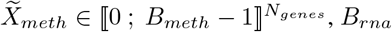 and *B*_*meth*_ being the number of bins respectively for gene expression and methylation. We fix the number of bins for both modalities at 64, and we present an ablation study that evaluates our tokenization strategy in Appendix A.6.

### 4.2. A long-range model architecture for bimodal representation learning

In order to learn representations of bulk RNA-seq and DNA methylation, we propose a model that combines both convolution and transformer blocks. Inspired by Avsec et al. (2021); Linder et al. (2025); Joshi et al. (2025) to handle long-range genomic dependencies, this architecture allows us to cope with the high dimensionality of the two omics that we consider, each corresponding to a sequence of length *N*_*genes*_. More precisely, a first bimodal embedding is obtained by passing each omic token through classic embedding layers and summing them along with a gene embedding vector. As gene expression and DNA methylation are permutation invariant, this gene embedding acts as a positional encoding and is randomly initialized. Prior to processing by a transformer model composed of multi-head attention layers (Vaswani et al., 2017), the embedding is downsampled by a convolutional tower. This downscaling allows the transformer block to act on a compressed embedding vector to significantly reduce the computational cost and time. While the convolutional architecture may be counterintuitive for unordered data, it acts as an efficient mechanism for dimensionality reduction. The original sequence length is restored using a deconvolutional tower with residual connections flowing from the downsampling blocks. Separate language modeling heads predict binned gene expressions and methylation. This architecture is summarized in Figure 1 (with a zoom in on the transformer part in Appendix A.2). We also compare *MOJO* to a variant that retains only its transformer component, which we refer to as *MOJO-Transformer*.

**Figure 1.**
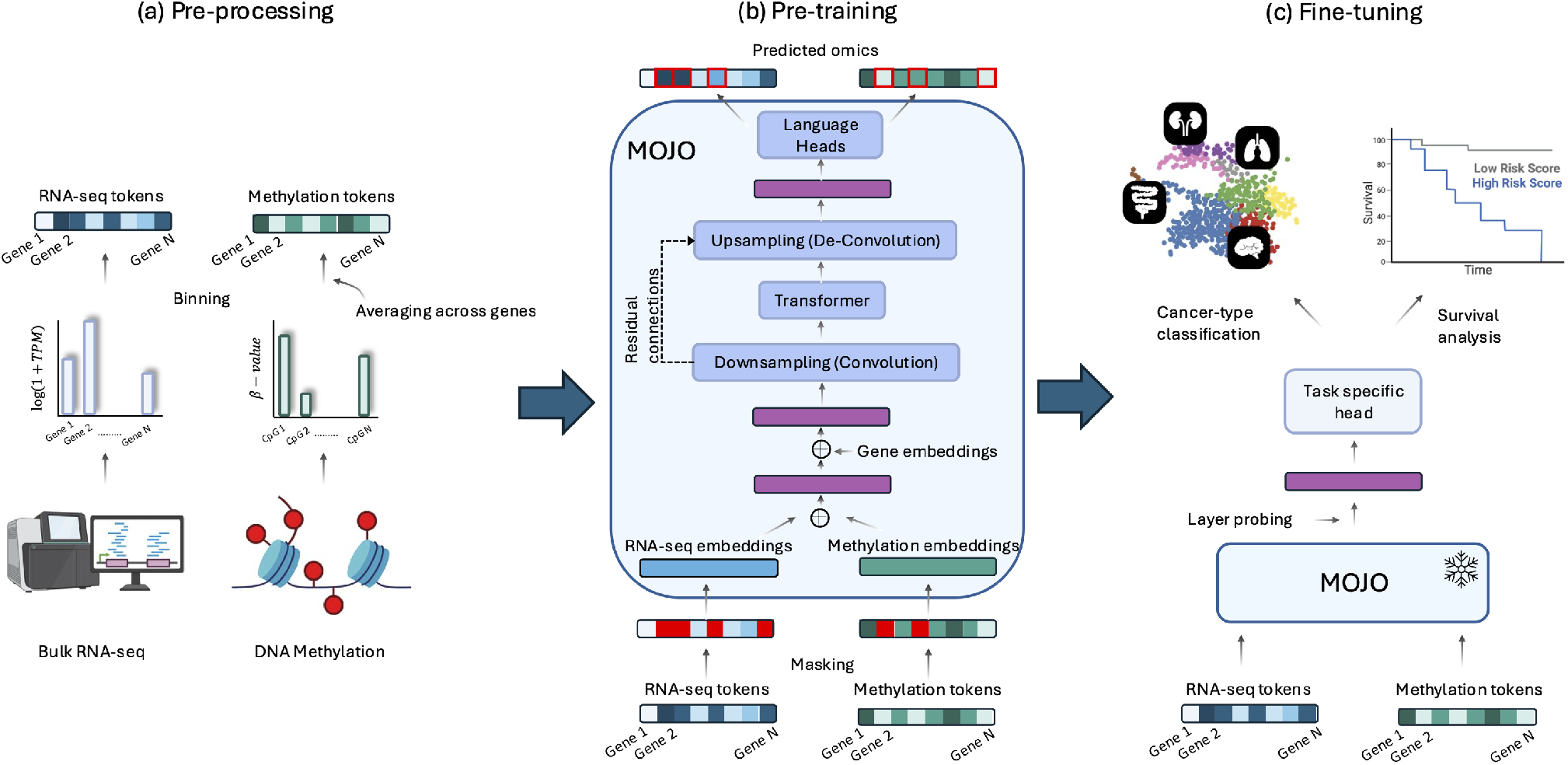
MOJO pipeline. (a) Each modality is first tokenized using linear binning. (b) *MOJO*, whose core architecture is composed of a mix of convolution and attention operations, is firstly pre-trained through bimodal masked language modeling. (c) Embeddings are probed from *MOJO* to fine-tune a task-specific head tailored for cancer-type classification or survival analysis.

### 4.3. Pre-training: bimodal masked language modeling

#### Self-supervision loss

Our model is pre-trained through self-supervision using multimodal masked language modeling. Although this framework may be applied to more than two modalities that can be processed as a sequence of tokens, we present it in the bimodal case where the set of modalities 𝕄 = {*rna, meth*}. We adopt standard parameters for masked language modeling (Devlin et al.,2019): for each sequence, 15% of the tokens are corrupted to train the model. Among these corrupted tokens, 80% are replaced with a special `<MASK>` token, 10% are substituted with random tokens, and the remaining 10% are left unchanged, but still contribute to the loss. The final heads of our model provide a set of probability distributions 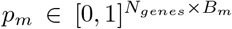 for *m* ∈ 𝕄. Denoting ℳ_*m*_, for *m* ∈ 𝕄, the set of masked token indices for that modality, the following multimodal negative log-likelihood is then optimized: 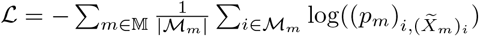

#### Experiment

Our model is pre-trained using the TCGA dataset from 33 cohorts, resulting in 9,252 pairs of bulk RNA-seq and methylation, 5% being kept for testing. We selected *N*_*genes*_ = 17,116 genes by using the same set of genes as Gélard et al. (2025), from which genes with no methylation data were removed. The model is trained on a total of 192 billion tokens using the Adam (Kingma & Ba, 2014) optimizer with gradient accumulation to reach an effective batch of 3 × 10^6^ tokens, on a TPU v4-8. The complete set of hyperparameters can be found in Appendix A.1, as well as pre-training learning curves in Appendix A.4.

## 5. Evaluation downstream tasks

The representations learned by *MOJO* are evaluated using a panel of downstream tasks ranging from supervised classification, survival analysis, and zero-shot classification, to clustering. We compare our method to unimodal (only RNA-seq or DNA methylation) and bimodal models:

### BulkRNABert (Gélard et al., 2025)

A transformer-based model pretrained on bulk RNA-seq using masked language modeling. Embeddings are extracted from the last self-attention layer, and the mean embedding over the sequence is used as input for downstream tasks. The tokenization of the RNA-seq data is the same between *MOJO* and *BulkRNABert, i*.*e*., the same *B*_*rna*_ has been used.

### scGPT (Cui et al., 2024)

We benchmark *scGPT* by adapting it from its original single-cell context to serve as a bulk RNA-seq encoder. This is achieved by applying the same parameter-efficient fine-tuning procedure used for *BulkRNABert*, effectively repurposing scGPT for bulk data across the different tasks.

### MethFormer

We develop the equivalent of *BulkRNABert* for DNA methylation (averaged per gene) and we will refer to it in the results as *MethFormer*. Similarly, the same value of *B*_*meth*_ is maintained to allow for fair comparison. This model differs from *MethylBERT* (Jeong et al., 2025) which uses read-level methylome and not the 450k microarrays we are interested in.

### Late integration

Bimodal integration resulting from the fusion of embeddings extracted from unimodal models. More precisely, we will refer to *Late integration (concatenation)* as the concatenation of the embeddings from *BulkRNABert* (for RNA-seq) and *MethFormer* (for methylation) which have been pre-trained beforehand. *Late integration (cross-attention)* corresponds to an integration of the two embeddings with a two-step cross-attention followed by a concatenation, allowing for interaction between the two modalities. The different cross-attention modules are only trained when fitting the downstream tasks. An illustration of the late integration is provided in Appendix B. Finally, we employ the integration strategy from **MultiSurv** (Vale-Silva & Rohr, 2021), where multimodal representations are aggregated by computing the maximum over features from unimodal models. We apply this aggregation method across all downstream tasks, including those beyond survival analysis, but retain the identifier *MultiSurv* to refer to this specific integration technique.

### CustOmics (Benkirane et al., 2023)

A multi-omics model based on Variational Auto-Encoders and tailored for cancer-type classification and survival analysis. Although it can handle up to three modalities (bulk RNA-seq, DNA methylation, and Copy Number Variation), we are here interested in its version that can perform the downstream tasks in the bimodal setting. Two models are considered: *CustOmics (end-to-end)* which trains the VAEs and the task heads jointly, and *CustOmics (probing)* that first learns the unsupervised representation with VAEs and then uses the encoded features as input to task heads.

### Multi-Omics Factor Analysis (*MOFA*) (Argelaguet et al., 2018)

An unsupervised machine learning method designed to integrate multi-omics data by inferring a set of low-dimensional hidden factors, which can be seen as a multiomics extension to PCA. *MOFA* factors are then used as input either to a Support Vector Machine (Cortes & Vapnik, 1995) for cancer-type classification or a Cox proportional model (Cox, 1972) for survival analysis.

### SeNMo (Waqas et al., 2024)

A self-normalizing MLP for multi-omics data integration, using ELU activations with *AlphaDropout* (Klambauer et al., 2017). We adapt *SeNMo* to operate on the concatenation of bulk RNA-seq and DNA methylation features, retrained on the same TCGA splits as MOJO.

### GAT

Inspired by MOGAT (Tanvir et al., 2024), this Graph Attention Network builds a *k*-nearest-neighbour patient-similarity graph over the dataset. Modality-specific linear projections compress RNA-seq and methylation features independently before fusion, and two GAT layers aggregate neighbourhood information to produce patient embeddings. We adapt it to operate on bulk RNA-seq and DNA methylation data exclusively.

An exhaustive benchmark of other feature extraction and multi-omics integration models is included in Appendix C.

### 5.1. Cancer-type classification

#### Methodology

*MOJO* is first evaluated on the supervised task of cancer-type classification. The pan-cancer TCGA dataset is divided into 33 cohorts that make up the labels for the classification task. The last attention layer of the transformer component of *MOJO* is probed to obtain the embedding used for classification. After being downsampled by the convolutional layers, the embedding lies in 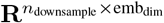, with emb_dim_ = 512 and *n*_downsample_ = 67 (resulting from 8 successive downsampling operations by a factor of 2 from an initial sequence length of *N*_genes_ = 17,116, padded to the next power of 2, 17,152). Then a mean embedding in 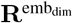 is obtained by averaging over the sequence dimension. This embedding serves as the input to a Multi-Layer Perceptron (MLP) of two hidden layers (respectively of size 256 and 128) that outputs logits for cancer-type prediction. Additionally, we apply dropout (Srivastava et al., 2014) and layer normalization (Ba et al., 2016). *BulkRNABert, MethFormer*, and *MOJO* are further fine-tuned in addition to training the MLPs using the parameter-efficient method *IA*^3^ (Liu et al., 2022), which introduces low-dimensional learnable parameters into the self-attention mechanisms and feed-forward networks. For *MOJO*, we also adapt this principle to the convolutional layers by performing a point-wise multiplication between the output of each convolutional operation and a learnable vector of the same dimension.

#### Results

Cancer-type classification results on the pan-cancer TCGA dataset are presented in Table 1. For this task, the dataset has been split into 80% for training and 20% for testing (stratified by cohort), repeating the split for 5 different seeds. We report the average and standard deviation over these 5 seeds for the different metrics. We will be using both macro *F*_1_ and weighted *F*_1_ scores to avoid any bias due to class imbalance (label distribution is provided in Appendix C.1). *MOJO* provides state-of-the-art results when considering the two modalities, with better performance than its pure Transformer equivalent (*MOJO-Transformer*), *CustOmics* and late integration. For the latter, the *cross-attention* version performs similarly to its *concatenation* counterpart. Our joint modeling of RNA-seq and methylation with *MOJO* outperforms the corresponding unimodal transformer-based models (*BulkRNABert* and *MethFormer*). Moreover, when only probing the last attention layer and fitting an SVM (*MOJO (probing)* in the table), our model shows a clear performance increase in comparison with *CustOmics (probing)*, highlighting that representations from masked language modeling exhibit stronger predictive capacity. Finally, we show that our bimodal masked language modeling pre-training produces a notable performance gain compared to a model trained from scratch (*MOJO (no pre-training)* in the table).

**Table 1.**
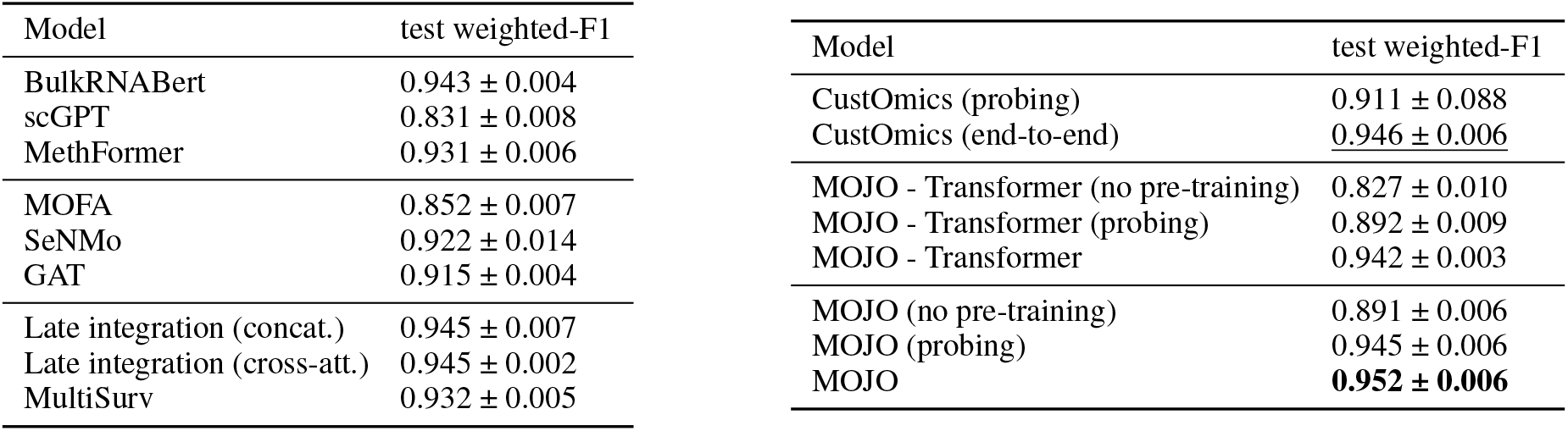
Cancer type classification. **Bold**: best; <u>underline</u>: second-best.

#### Training times

We additionally report in Table 2 the time required by different models (*BulkRNABert, Late integration (cross-attention), Late integration (concatenation), MOJO*, and *MOJO - Transformer*) to perform a full update step (forward and backward pass) when training a pan-cancer classification model. While supporting substantially larger batch sizes compared to purely transformer-based models or late integration mechanisms, MOJO achieves approximately a 100*×* speedup over other benchmarked models, and a 300*×* speedup over its transformer-based counterpart. This highlights the computational efficiency of our hybrid architecture that combines convolutional and transformer layers, offering a more scalable alternative to fully transformer-based approaches.

**Table 2.** Average time per update step (forward + backward pass) during training of classification models on a TPU v4-8. All models are evaluated with an effective batch size of 64, achieved via gradient accumulation when necessary. For each model, we additionally report the maximum batch size supported by the model. As in classification benchmarks, parameter-efficient fine-tuning is applied to *MOJO, MOJO - Transformer*, and *BulkRNABert*.

| Model | Update time (seconds) | Maximum batch size |
| --- | --- | --- |
| BulkRNABert | 4.462 ± 0.006 | 8 |
| Late integration (cross-attention) | 5.819 ± 0.006 | 4 |
| Late integration (concatenation) | 2.205 ± 0.004 | 16 |
| MOJO - Transformer | 17.987 ± 0.001 | 4 |
| MOJO | **0.059 ± 0.009** | **1,024** |

### 5.2. Survival Analysis

#### Methodology

We then evaluate omics embeddings on a pan-cancer survival task, also known as time-to-event prediction. This task involves predicting the survival time 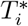 for individuals with cancer, specifically the time from diagnosis until death. A key challenge in survival analysis is right censoring, where the observed time 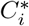 might be shorter than the actual survival time 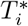 due to factors like the end of a study or loss of patient contact. Consequently, the true target time used by the model, *T*_*i*_, is defined as the minimum of the actual survival time and the censoring time 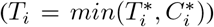. One defines as well 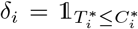 (so *δ*_*i*_ = 1 if the event occurred (death), otherwise *δ*_*i*_ = 0), thus constituting a dataset 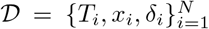, with *x*_*i*_ the covariates (RNA-seq and/or methylation embeddings in our study). To ensure robust evaluation, data splits for this task were stratified based on the (cohort, event) pair, guaranteeing that each split maintains a consistent distribution of cohorts and observed mortality events. A widely used method to tackle such time-to-event problems is the Cox proportional hazards model (Cox, 1972). This semi-parametric model focuses on modeling the hazard function *λ*(*t* |*x*), which represents the instantaneous rate of an event at time *t* given covariates *x*. A Cox model expresses *λ* as 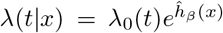, with *β* a vector of parameters (so in our case *x*,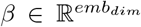), *λ*_0_(*t*) the hazard baseline, and in the Cox model, *ĥ*_*β*_ (*x*) = *β*^*T*^ *x*. More recent works (like *DeepSurv*, or *Cox-nnet* (Katzman et al., 2018; Ching et al., 2018) relax the linear combination of features to replace *ĥ*_*β*_ (*x*) with the output of a neural network. For the survival task, we extract the embeddings from *BulkRN-ABert, MethFormer*, and *MOJO* in the same way as for the cancer-type classification task and train a similar MLP architecture on top. We do not consider the cross-attention version of the late integration here as the Cox loss requires working with a sufficiently large batch size, rendering the computation of cross-attentions (given the sequence length of *N*_*genes*_ = 17, 116) computationally prohibitive so that the computation of cross-attentions. Similarly, we do not apply *IA*^3^ and only consider the probing experiment for the survival task (as fine-tuning with *IA*^3^ yielded negligible improvements over probing for this task, see Appendix C.3).

#### Results

Survival results on the pan-cancer TCGA dataset are presented in Table 3. The same split strategy as for the classification task is used. Two different evaluation metrics based on Harrell’s C-index (Harrell et al., 1982) are reported. First, a C-index is computed on the whole test set (all cohorts) referred to as “C-index”. However, in order to make sure that a pan-cancer model is able to predict survival within cohorts correctly, and not just to differentiate survival chances between cancer types, a “Weighted C-index” is also reported. This corresponds to a weighted sum of the C-indexes computed per cohort on the pan-cancer test set, with weights corresponding to the number of samples of each cohort in the test set. As for the classification task, *MOJO* exhibits higher performance than the unimodal transformer-based models and the late integration. *MOJO* (52M parameters) outperforms *CustOmics* (46M) with a significant gain, confirming that architectural design, specifically the hybrid convolutional-transformer pre-training, rather than model capacity alone drives performance. In an additional experiment, *MOJO*’s performance is matched by an end-to-end version of *CustOmics* (weighted C-index of 0.669 ± 0.004). This need for end-to-end training compared to simple probing highlights the strength of *MOJO*’s learned representations. *MOJO* also outperforms *SeNMo* and *GAT*; notably, the *GAT* model achieves a competitive global C-index but the lowest weighted C-index (0.619), revealing that inter-patient graph comparisons benefit cross-cohort ranking but fail to capture within-cohort survival variation. Kaplan-Meier curves are provided in Appendix C.5, showing better patient stratification with *MOJO*.

**Table 3.** Pan-cancer survival analysis.

| Model | C-index | Weighted C-index |
| --- | --- | --- |
| BulkRNA-Bert | $0.749 \pm 0.003$ | $0.654 \pm 0.014$ |
| scGPT | $0.720 \pm 0.005$ | $0.604 \pm 0.020$ |
| MethFormer | $0.736 \pm 0.006$ | $0.622 \pm 0.014$ |
| MOFA | $0.648 \pm 0.037$ | $0.601 \pm 0.022$ |
| CustOmics | $0.686 \pm 0.018$ | $0.639 \pm 0.099$ |
| Late int. (concat.) | $0.756 \pm 0.004$ | $0.653 \pm 0.011$ |
| MultiSurv | $0.742 \pm 0.002$ | $0.636 \pm 0.011$ |
| SeNMo | $0.745 \pm 0.004$ | $0.640 \pm 0.017$ |
| GAT | $0.757 \pm 0.007$ | $0.619 \pm 0.013$ |
| MOJO-Transformer | $0.757 \pm 0.006$ | $0.657 \pm 0.007$ |
| MOJO | **$0.771 \pm 0.006$** | **$0.670 \pm 0.009$** |

### 5.3. Zero-shot pan-cancer and breast cancer sub-typing and clustering

#### Methodology

To evaluate *MOJO*’s learned embeddings in a fully unsupervised manner, we assess their zero-shot classification and clustering capabilities on PAM50 breast cancer sub-typing (Luminal A, Luminal B, Basal, and HER2) (Parker et al., 2009) and the Pan-cancer dataset from section 5.1. First, zero-shot classification uses a *k*-nearest neighbors model (*k* = 5), evaluated by accuracy using 5-fold stratified cross-validation, to assess embedding quality without fine-tuning, inspired by Joshi et al. (2025). Second, Leiden clustering (Traag et al., 2019) is performed in the embedding space, with Normalized Mutual Information (NMI) and Adjusted Rand Index (ARI) as metrics. We primarily compare the effectiveness of *MOJO*’s joint modeling against late integration (concatenation) embeddings for bimodal data.

#### Results

Zero-shot classification and clustering results are shown in Table 4, showing better performance when using *MOJO* embedding than late integration. A comparison with *CustOmics* is also added in Appendix C.2, with lower performance than *MOJO*. Figure 2 presents t-SNE plots of both embeddings in the pan-cancer setting, demonstrating that MOJO embeddings achieve superior cohort separation.

**Table 4.** Zero-shot classification and clustering results on pan-cancer and PAM50 tasks. (Acc. = Accuracy, NMI = Normalized Mutual Information, ARI = Adjusted Rand Index).

| Task | Metric | MOJO | Late integration |
| --- | --- | --- | --- |
| PAM50 | Acc. | **0.777** | 0.763 |
|  | NMI | **0.345** | 0.291 |
|  | ARI | **0.213** | 0.154 |
| Pan-cancer | Acc. | **0.928** | 0.870 |
|  | NMI | **0.862** | 0.771 |
|  | ARI | **0.756** | 0.620 |

**Figure 2.**
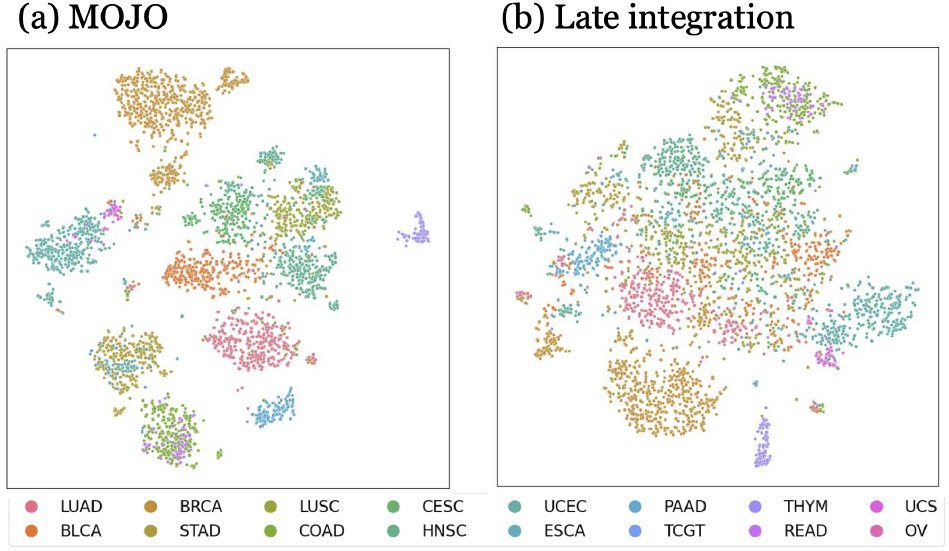
t-SNE representation of *MOJO* and *Late integration* embeddings, colored by cancer-type on a subset of cohorts. *MOJO*’s embeddings visually exhibit better cohort separation capacity compared to *Late Integration* ones, corroborating quantitative results from Table 4. Full pan-cancer t-SNE plot is provided in Appendix C.4.

### 5.4. MOJO encodes known biological pathways

While *MOJO*’s representations seem to be powerful for predictive tasks like cancer-type classification and survival analysis, interpreting them is also a critical step. To this end, we focus on a classification model based on *MOJO*’s embeddings (same setting as per Section 5.1) on the PAM50 BRCA sub-typing task (presented in section 5.3). Our classification model maps *MOJO’s* embeddings of 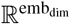 to logits in 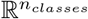 with here *n*_*classes*_ = 4. We compute Shapley (SHAP) Values (Lundberg & Lee, 2017) for this model and focus on a particular subtype. We identify the most important dimension as the one that maximizes the mean absolute SHAP value. However, SHAP values only inform us about the contributions of the embedding dimensions to the predictions, but without indicating which genes drove the activation of these dimensions. To return to the gene level, for each gene, we compute the Spearman correlation between the value of the embedding at that dimension and the bulk RNA-seq value. This yields a ranked list of genes that is used as input to a pre-ranked Gene Set Enrichment Analysis (GSEA, Subramanian et al. (2005)). We extend this approach to multiple dimensions by considering the average of the Fisher z-transformations of the correlation obtained for the *top k* best dimensions (as per *Stouffer*’s Z-score method (Stouffer et al., 1949)). Taking the example of *Luminal B* sub-type, Figure 3 shows that known pathways are correctly enriched (like *Protein Secretion* or *Estrogen Response*) (Li et al., 2016; Tran & Bedard, 2011; Fertig et al., 2015), showing that the dimensions that contribute the most to classifying this subtype encode known biological information (*k* = 10 has been used here). Extended results on all BRCA subtypes can be found in Appendix F.

**Figure 3.**
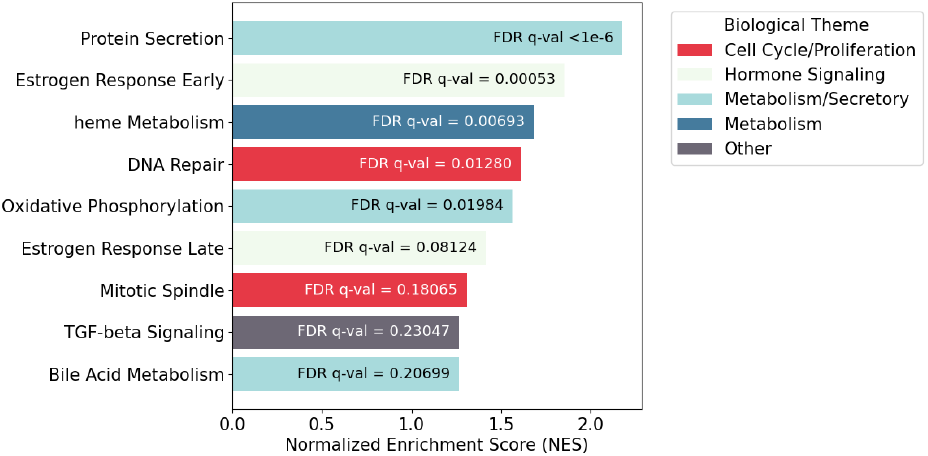
Cancer-relevant enriched pathways for *LuminalB* subtype using pre-rank GSEA from correlations with most informative *MOJO*’s dimensions. Only pathways with positive enrichment scores are represented. (*FDR*: False Detection Rate).

## 6. External validation and platform generalization

### 6.1. Cross-platform transferability to legacy 27k arrays

*MOJO* aggregates CpG probes into gene-level methylation features, making it inherently array-agnostic. To test transferability to legacy 27k platforms, we generated an in-silico 27k dataset by filtering the TCGA 450k data to the 27k microarray probe IDs, maintaining exact patients for a 1-to-1 comparison. Since 27k arrays are heavily promoter-biased, the resulting gene-level values differ substantially from their 450k counterparts (mean shift of − 0.115), inducing a severe distribution shift. Under zero-shot evaluation, baseline models collapse: *CustOmics*’ macro-*F*_1_ drops from 0.922 to 0.226 and survival C-index from 0.686 to 0.509. In contrast, zero-shot *MOJO* retains robust performance (C-index: 0.717; macro-*F*_1_: 0.734), even outperforming a *CustOmics* model fine-tuned on 27k arrays on the survival task. Furthermore, per-gene z-score rescaling (still without any fine-tuning) recovers near full-450k performance (C-index: 0.765; macro-*F*_1_: 0.909). Detailed methodology and extended metrics are provided in Appendix G.

### 6.2. Generalization to external held-out cohorts

We assessed *MOJO*’s generalization on two held-out external datasets. On the TARGET cohort, while survival performance is on par with *BulkRNABert* (C-index 0.601 ± 0.023), suggesting methylation provides overlapping rather than additive prognostic information in this pediatric setting, *MOJO*’s bimodal representations demonstrate clear advantages in zero-shot clustering, with substantially higher NMI (0.949 vs. 0.794) and ARI (0.946 vs. 0.706) over late integration. On the ICGC cohort of 405 adult donors across four cancer types, *MOJO* achieves a superior C-index of 0.823 ± 0.019 (Table 5), outperforming both *BulkRNABert* (0.805 ± 0.036) and *MethFormer* (0.810 ± 0.027), confirming that MOJO’s bimodal embeddings generalize to entirely unseen cohorts. Complete dataset details and extended evaluation are provided in Appendix E.

**Table 5.** Survival on the external ICGC cohort.

| Model | C-index | Weighted C-index |
| --- | --- | --- |
| BulkRNABert | $0.805 \pm 0.036$ | $0.610 \pm 0.087$ |
| MethFormer | $0.810 \pm 0.027$ | $0.628 \pm 0.062$ |
| MOJO | **$0.823 \pm 0.019$** | **$0.643 \pm 0.032$** |

## 7. Missing modalities

While *MOJO*’s joint modeling of two modalities improves downstream task performance, clinical applications may face missing modalities, including the potential for a modality to be entirely unavailable. Being trained by masked language modeling, *MOJO* naturally accepts missing RNA-seq or methylation information for a given subset of genes by attributing a special <MASK> token to these genes. One can naturally extend this procedure by assigning a sequence entirely composed of <MASK> tokens to a missing modality, making *MOJO* inherently capable of handling missing modalities. However, as during pre-training only a fraction of each modality is masked to account for the masked language modeling loss, the absence of a whole modality is never encountered by the model. To this end, we conducted an additional pre-training of *MOJO* by incorporating samples from the TCGA dataset that are missing one of the two considered modalities, thus extending the initial pre-training dataset composed of 9,252 pairs (*X*_*rna*_, *X*_*meth*_) with 2,022 pairs (*X*_*rna*_, *None*) and 560 pairs (*None, X*_*meth*_) with *None* indicating a missing modality. We will refer to this model as *MOJO-Extended-Pretraining*. Two settings thus arise: **(S1)** one modality is either missing during model fine-tuning and we want to assess the utility of *MOJO*’s embeddings in this situation, or **(S2)** it is missing only at test-time while the downstream model has been trained with full bimodal input.

### 7.1. Missing modalities for fine-tuning (S1)

In the context of Ovarian (OV) cancer subtyping in TCGA (4 classes: differentiated, immunoreactive, mesenchymal, and proliferative (Verhaak et al., 2012)), only RNA-seq samples are available. A bimodal pre-trained *MOJO* model is thus fine-tuned on this task with (*X*_*rna*_, *None*) as input and achieves superior performance than *BulkRNABert* while being faster to train (Figure 4). Using *MOJO-Extended-Pretraining* instead of *MOJO* further improves the performance in OV subtyping. In this context, we prove that *MOJO*, while initially being a bimodal model, can be used as an embedding model for RNA-seq only, and provide better performance than a specific unimodal model.

**Figure 4.**
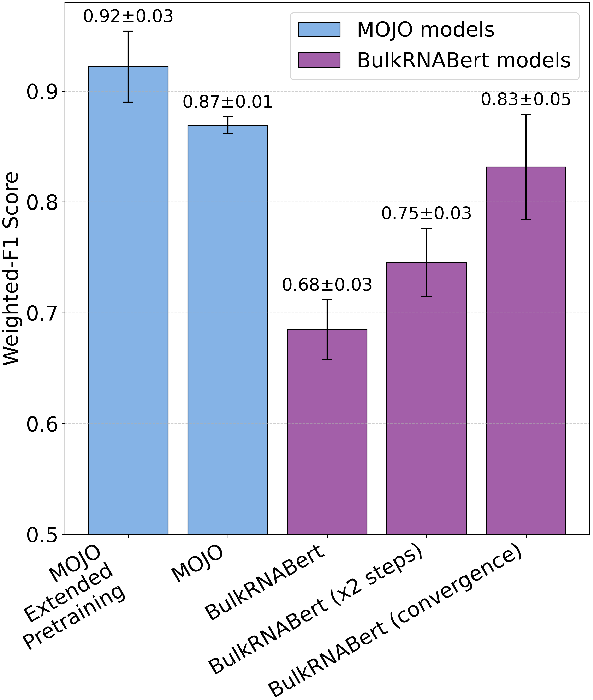
Ovarian cancer sub-typing: MOJO outperforms *BulkRNABert* while being faster to fine-tune. (“*BulkRNABert* models” bars from left to right: same fine-tuning budget as *MOJO*, ×2 fine-tuning steps, and until convergence). *MOJO-Extended-Pretraining* refers to an extension of the pre-training dataset with samples with missing modalities.

### 7.2. Missing modalities at test-time: Mutual Information loss (S2)

#### Motivation

Our goal is to create bimodal models for tasks like the pan-cancer classification task (for which both modalities are available during fine-tuning) that maintain performance comparable to unimodal models when a modality is absent at test time. Our framework begins with training a bimodal downstream model fine-tuned using complete (*X*_*rna*_, *X*_*meth*_) pairs from a pre-trained *MOJO* (per section 5.1). Our contribution involves modifying this fine-tuning for missing modalities.

#### Methodology

At test time, we compute metrics under two settings: with the complete test set, and by simulating the absence of either RNA-seq or methylation by dropping it from x% of test pairs (up to x=100% for removal in all samples), all without further model fine-tuning. We only fine-tune and evaluate the downstream model with samples that have both modalities, thus allowing us to get a fair comparison between all the models. Since the dataset is fixed, we must modify the fine-tuning procedure to maintain performance despite the complete absence of a modality at test time. To this end, we add mutual information as an auxiliary loss paired with classic cross-entropy for the pan-cancer classification task. This quantity is used in Ramazanova et al. (2025) as a test-time adaptation technique of an audio/vision model to handle missing modalities. We adapt it to be directly incorporated during the fine-tuning phase of the model to avoid any modification of the model at test time, thus saving computational time. Following the notation from Ramazanova et al. (2025), we denote *f*_*θ*_(*x*; *m*) the output of the classification model for a given input *x* when modalities *m* are present, with *m* ∈ 𝒟_*modality*_ = {*rna* + *meth, rna, meth*}. One would require *f*_*θ*_ to provide the same prediction regardless of the provided modality, thus satisfying the following equality: *f*_*θ*_(*x*; *rna* + *meth*) = *f*_*θ*_(*x*; *rna*) = *f*_*θ*_(*x*; *meth*). One can satisfy such a constraint by minimizing the following loss: 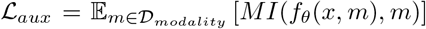 with *MI*(*X, Y*) = *D*_*KL*_[*P*_(*X,Y*)_ || *P*_*X*_ ⊗ *P*_*Y*_] corresponding to the mutual information (Shannon, 1948) between two random variables *X* and *Y*. The mutual information is equal to 0 when the two random variables are independent; thus, minimizing this quantity as an auxiliary loss should guide the model towards providing the same output independently of the given modality. Therefore, the following loss is now optimized: ℒ = ℒ_*task*_ +*λ*\* ℒ_*aux*_, with ℒ_*task*_ corresponding here to the cross-entropy for the cancer-type classification task. We detail in the algorithm of Appendix H the procedure used to compute this loss for a single example.

#### Results

Results from the missing modalities experiments are shown in Figure 5 when RNA-seq is dropped (similar results when dropping DNA methylation in Appendix H). The initial *MOJO* model achieved a test weighted-*F*_1_ of 0.952. Dropping all RNA-seq from the test set (*x* = 100%) significantly decreases performance to 0.538. Adding mutual information during fine-tuning (we used *λ* = 10 as the ratio of the cross-entropy and the mutual information computed after model initialization, see Appendix H.3 for an ablation study on this hyperparameter) maintained stable performance (0.949) with no modality drop. However, it corrected the performance decrease when RNA-seq was missing (from 0.538 to 0.916). These recovered scores approach those of unimodal models (*i*.*e, MethFormer* as methylation is the remaining modality). Extending pre-training with pairs missing one modality (*MOJO-Extended-Pretraining*) also narrowed the performance gap without the mutual information loss, and combining *MOJO-Extended-Pretraining* with the auxiliary mutual information loss yielded performance similar to using the auxiliary loss alone.

**Figure 5.**
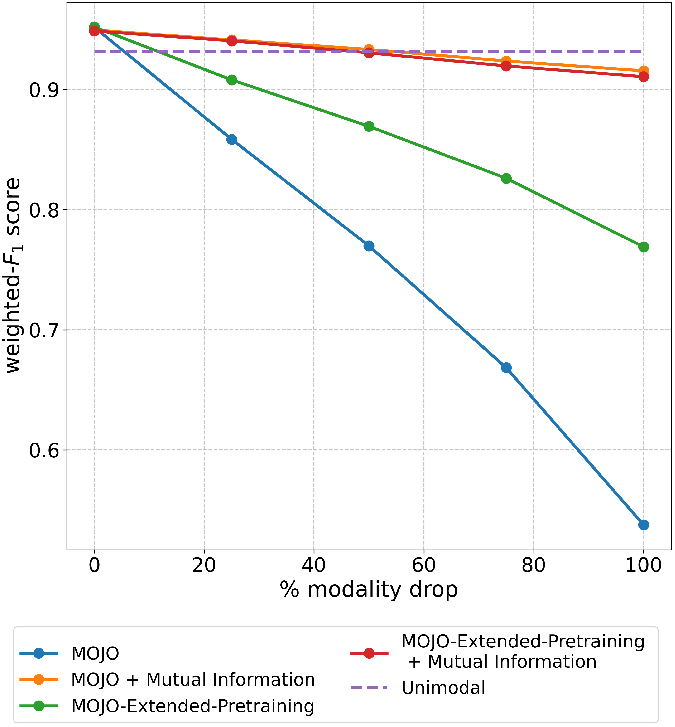
Performance when dropping RNA-seq. Test weighted-F1 score is reported as a function of the percentage of dropped RNA-seq samples in the test set. “*+ Mutual Information*” indicates the use of MI auxiliary loss during fine-tuning.

## 8. Conclusion

We introduced *MOJO*, a novel methodology for learning joint representations from multi-omics data, specifically bulk RNA-seq and DNA methylation. Pre-trained using self-supervised bimodal masked language modeling, *MOJO*’s architecture, combining convolutional and attention components, efficiently handles long gene sequences, outperforming purely transformer-based unimodal models. *MOJO*’s learned embeddings achieve state-of-the-art performance in supervised tasks like cancer-typing and time-to-event prediction on the TCGA dataset compared to unimodal models. In particular, the predictive capacity of *MOJO*’s representations is highlighted by the significant performance gain observed in the layer probing setup. Joint modeling’s advantage over late integration was demonstrated through zero-shot classification and clustering. Although *MOJO* inherently accepts missing modalities, we narrowed the performance gap with unimodal models which arises when a modality is missing by incorporating a mutual information-based loss during fine-tuning. Future work includes extending *MOJO* to more data types, relaxing modality alignment requirements, extending the mutual information approach to more modalities, and improving it to exactly match unimodal performance.

### Limitations

This work presents a robust framework for multi-omics integration and missing modality handling, improving the potential clinical utility of deep learning-based cancer prognosis. However, some limitations remain. First, *MOJO*’s gene-level methylation averaging acts as an information bottleneck, discarding potentially site-level biological variations. Second, while pre-trained on the entirety of the paired TCGA cohort, this dataset is small relative to other modality foundation models and inherently reflects TCGA’s specific population demographic biases. Deploying such prognostic models requires extensive prospective validation, as relying on these biases may lead to skewed clinical predictions across different populations.

## Impact Statement

This paper presents work towards an integration of multiomics data while handling missing modalities. This approach improves the clinical utility of deep learning-based cancer prognosis by enabling more reliable patient stratification.

## Acknowledgements

Research supported by Cloud TPUs from Google’s TPU Research Cloud (TRC).

## A. MOJO pre-training

### A.1. Hyperparameters

**Table 6.** MOJO model and pre-training hyperparameters.

| Model Hyperparameters |  |
| --- | --- |
| Number of downsamples | 8 |
| Kernel size | 5 |
| Embedding dimension | 512 |
| Number of transformer layers | 8 |
| Feed forward dimension | 1,024 |
| Number of attention heads | 16 |
| Training Hyperparameters |  |
| Batch size | 128 |
| Gradient accumulation | 4 |
| Learning rate | $5 \times 10^{-5}$ |
| Masking ratio | 15% |

### A.2. MOJO Transformer block

**Figure 6.**
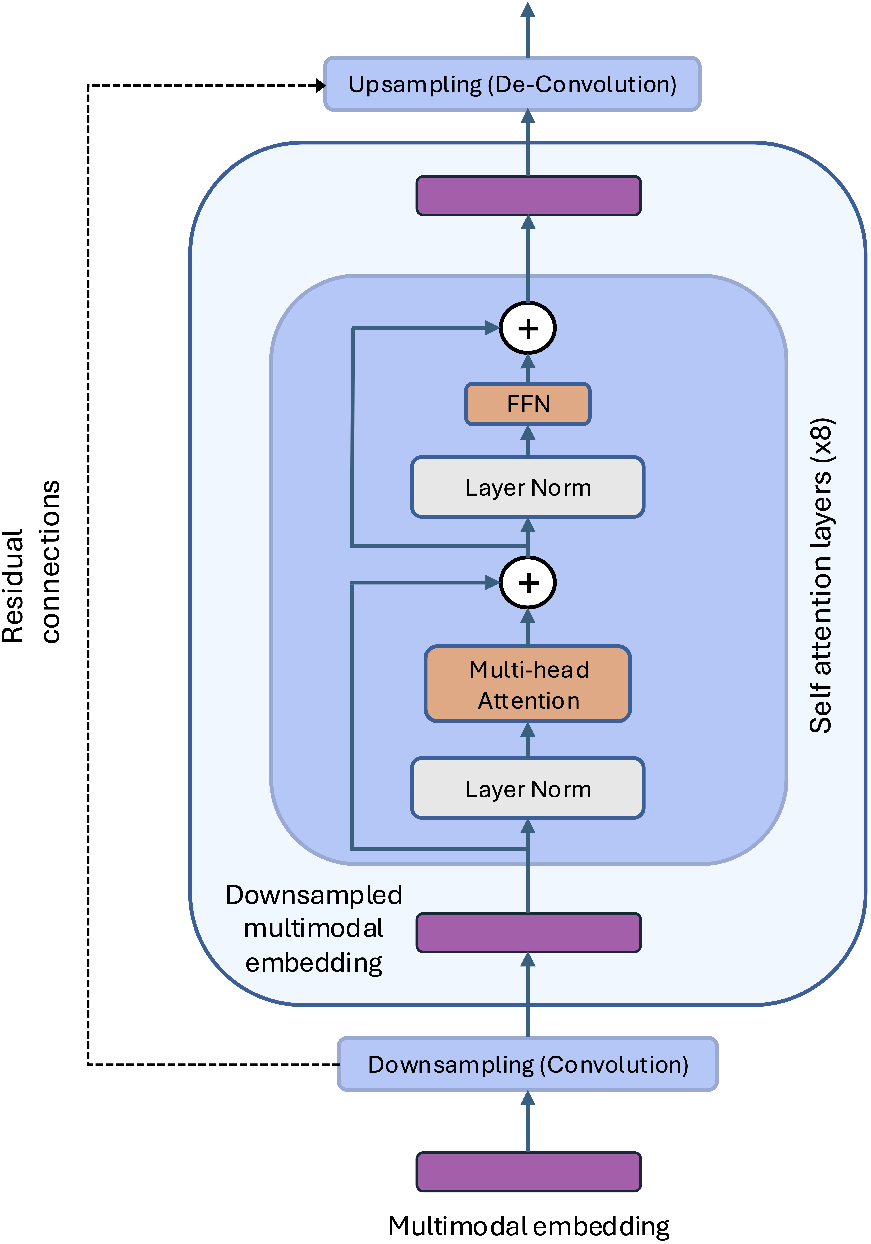
Zoom on the Transformer block of the *MOJO* architecture. Full architecture is provided in Figure 1.

### A.3. Pre-training dataset statistics and preprocessing

**Table 7.** Composition of the MOJO pre-training datasets.

| Model | Split | # Samples | Modality | Cohorts |
| --- | --- | --- | --- | --- |
| MOJO | Train | 8,789 (95%) | Bimodal (RNA-seq + Methylation) | 33 TCGA cohorts |
|  | Test | 463 (5%) | Bimodal (RNA-seq + Methylation) |  |
|  | **Total** | **9,252** |  |  |
| MOJO-EXTENDED | Train | 11,242 (95%) | Bimodal + RNA-seq only + Methylation only | 33 TCGA cohorts |
|  | Test | 592 (5%) | Bimodal + RNA-seq only + Methylation only |  |
|  | **Total** | **11,834** |  |  |

#### Preprocessing pipeline

RNA-seq values (TPM) were transformed as *x* ↣ log_10_(1 + *x*). For methylation, each sample’s 450k beta-values were mapped to gene-level averages by averaging all CpG sites annotated to that gene in the Illumina HumanMethylation450k BeadChip manifest (simple unweighted average, see Appendix D for the region-aware alternative). We retain *N*_*genes*_ = 17,116 genes present in both the RNA-seq and 450k annotations. Both modalities were tokenized using linear binning into 64 bins (RNA-seq: [0, log_10_(1 + max)]; methylation: [0, 1]). The vocabulary comprises 64 *×* 2 omic tokens plus special tokens (`<MASK>, <PAD>`).

### A.4. Pre-training learning curves

**Figure 7.**
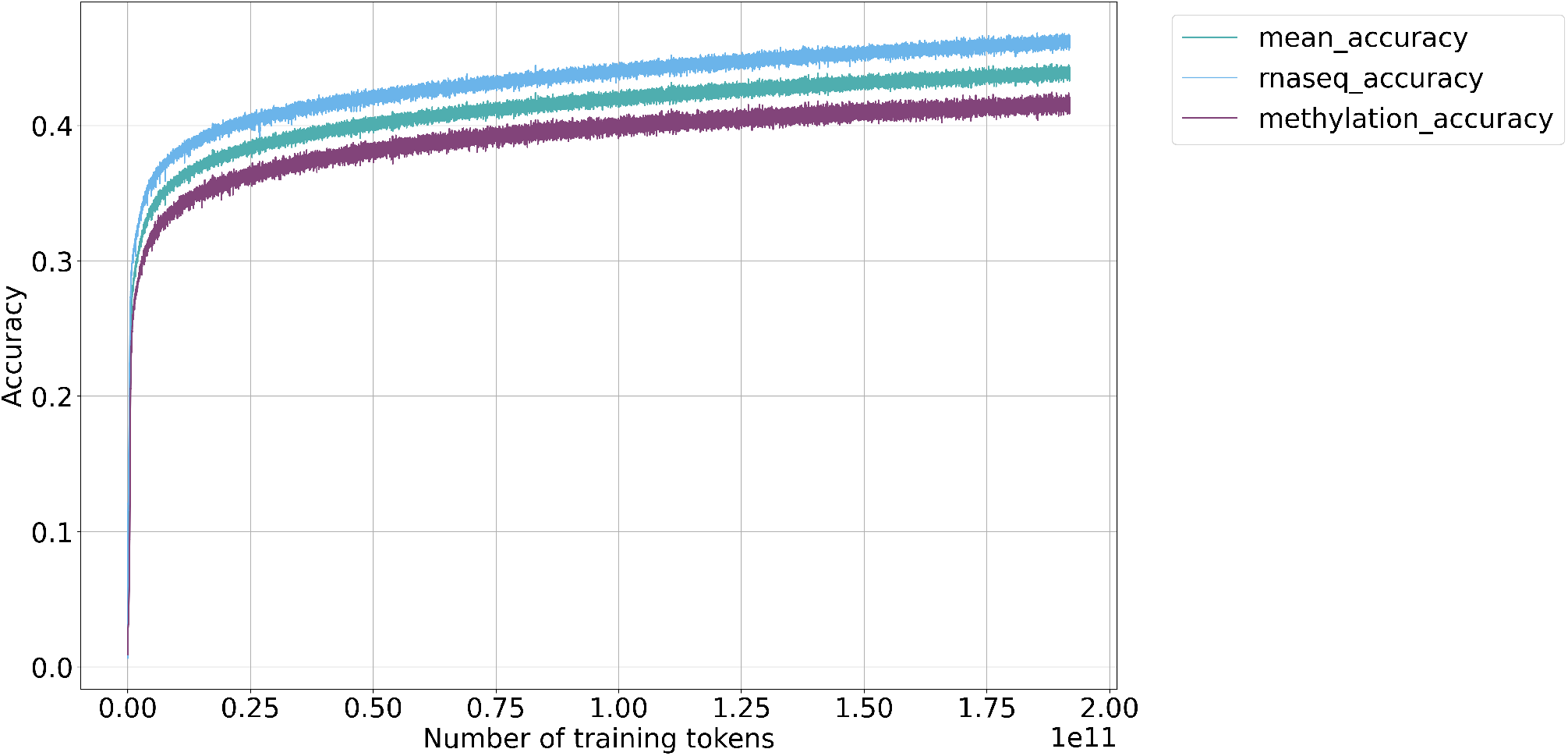
Bimodal masked language modeling pre-training curves of the *MOJO* architecture. The training reconstruction accuracy is represented for each omic separately as well as the average reconstruction accuracy among the different omics.

### A.5. Architectural justification: convolutions and input ordering

A potential concern regarding the application of convolution-based architectures, such as the U-Net, to bulk omics data is the lack of inherent spatial locality in the input. Unlike images, where pixel adjacency correlates with semantic structure, genes or CpG sites in a feature vector do not possess a natural grid-like order. However, our adoption of the U-Net backbone is motivated by computational efficiency and multi-scale representation rather than spatial inductive biases. Specifically, the U-Net facilitates hierarchical dimensionality reduction, essential for handling high-dimensional inputs (hundreds of thousands of features), and enables the aggregation of signals ranging from individual gene values to broad molecular patterns.

To empirically validate that the model’s performance does not rely on arbitrary input adjacency, we evaluated a variant, MOJO-ordered, where input features were sorted by genomic coordinates (chromosome and physical position). This ordering imposes a biological locality that a convolutional network could theoretically exploit.

As shown in Table 8, MOJO-ordered performs comparably to the standard model across both classification and survival tasks, with differences falling within the margin of error. These results confirm that the architectural benefits of MOJO stem from its capacity for hierarchical feature extraction in high-dimensional spaces, rather than local spatial correlations. Consequently, we retain the unordered design for its generality and simplicity.

**Table 8.** Impact of genomic ordering: performance comparison between the standard unordered MOJO and the genomically sorted MOJO-ordered variant. Results indicate that imposing biological order yields no statistically significant improvement, suggesting the architecture is effectively order-invariant in this context.

| Model | Cancer-type Classification |  | Survival Analysis |  |
| --- | --- | --- | --- | --- |
|  | Macro-F1 | Weighted-F1 | C-index | Weighted C-index |
| MOJO (Unordered) | 0.935 ± 0.007 | 0.952 ± 0.005 | 0.771 ± 0.006 | 0.670 ± 0.009 |
| MOJO-ORDERED | 0.936 ± 0.006 | 0.954 ± 0.002 | 0.775 ± 0.007 | 0.672 ± 0.019 |

### A.6. Tokenization sensitivity analysis

We analyzed the discretization strategy used to tokenize continuous bulk RNA-seq and DNA methylation values. All models were pre-trained from scratch with the corresponding binning scheme and fine-tuned on the same downstream tasks. Two strategies are compared: *linear* tokenization, which divides the value range into equally spaced bins, and *quantile* tokenization, which sets bin boundaries according to each gene’s empirical distribution so that each bin contains approximately the same number of samples.

**Table 9.** Sensitivity of *MOJO* to tokenization strategy and bin granularity. **Bold**: default configuration.

| Strategy | Bins | Cancer-type Classification |  | Survival Analysis |  |
| --- | --- | --- | --- | --- | --- |
|  |  | Macro-F1 | Weighted-F1 | C-index | Weighted C-index |
| Linear | 5 | 0.910 ± 0.007 | 0.937 ± 0.005 | 0.758 ± 0.009 | 0.650 ± 0.020 |
| Linear | 16 | 0.925 ± 0.005 | 0.948 ± 0.004 | 0.770 ± 0.007 | 0.669 ± 0.013 |
| Linear | 32 | 0.937 ± 0.006 | 0.953 ± 0.004 | 0.772 ± 0.005 | 0.674 ± 0.020 |
| **Linear** | **64 (default)** | **0.935 ± 0.007** | **0.952 ± 0.006** | **0.771 ± 0.006** | **0.670 ± 0.009** |
| Linear | 128 | 0.941 ± 0.006 | 0.954 ± 0.003 | 0.770 ± 0.007 | 0.664 ± 0.021 |
| Quantile | 64 | 0.934 ± 0.008 | 0.952 ± 0.006 | 0.771 ± 0.006 | 0.670 ± 0.008 |

Performance is stable for ≥ 32 bins, with 64 bins providing the best trade-off between capturing fine-grained biological variation and filtering technical noise. The coarser 5-bin scheme degrades macro-*F*_1_ by 2.7 points and C-index by 1.7 points, confirming that sufficient resolution is important for discriminating subtle expression states. Quantile-based tokenization achieves results equivalent to linear scaling, validating our default tokenization choice.

## B. Late integration

An illustration of the late integration mechanism is provided in Figure 8.

**Figure 8.**
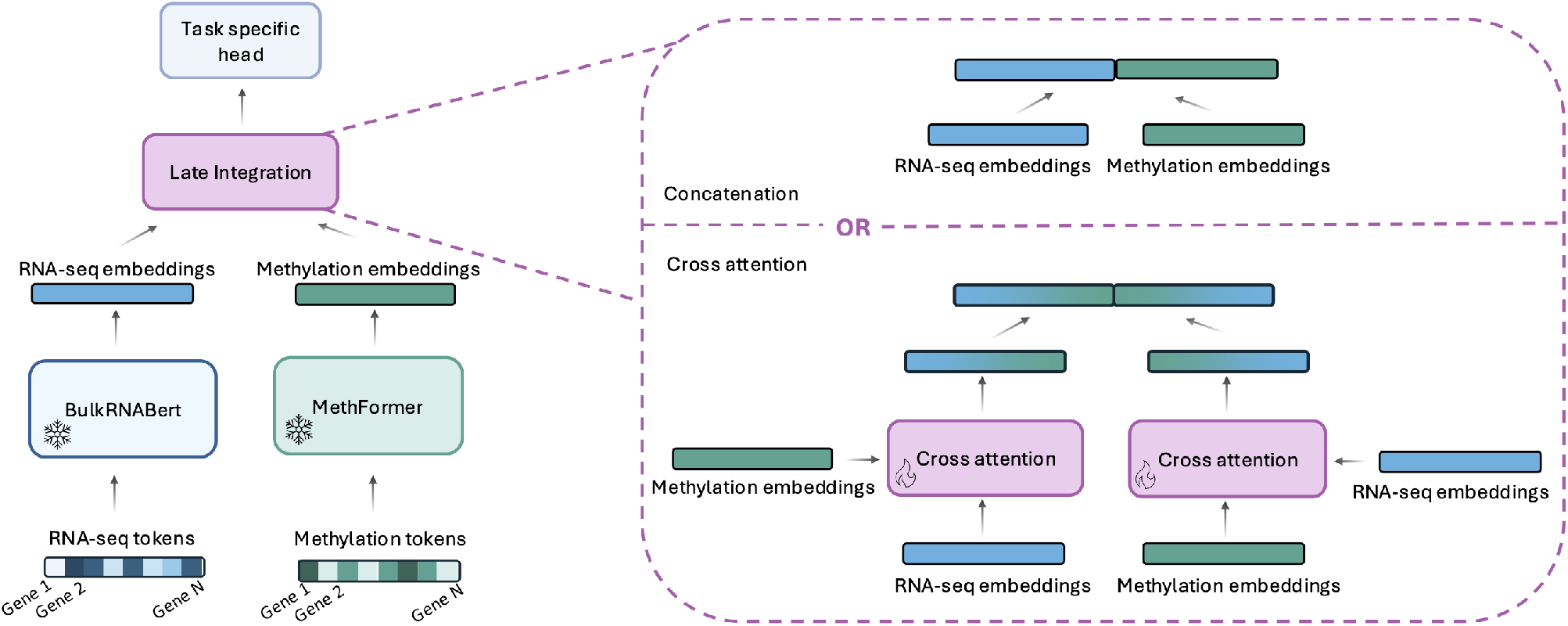
Late integration architecture. RNA-seq and methylation embeddings are obtained from pre-trained transformer based encoders (respectively *BulkRNABert* and *MethFormer*) and are fused either by concatenation or by a two-step cross-attention mechanism.

## C. Downstream tasks dataset and benchmarks

### C.1. Pan-cancer classification dataset

**Figure 9.**
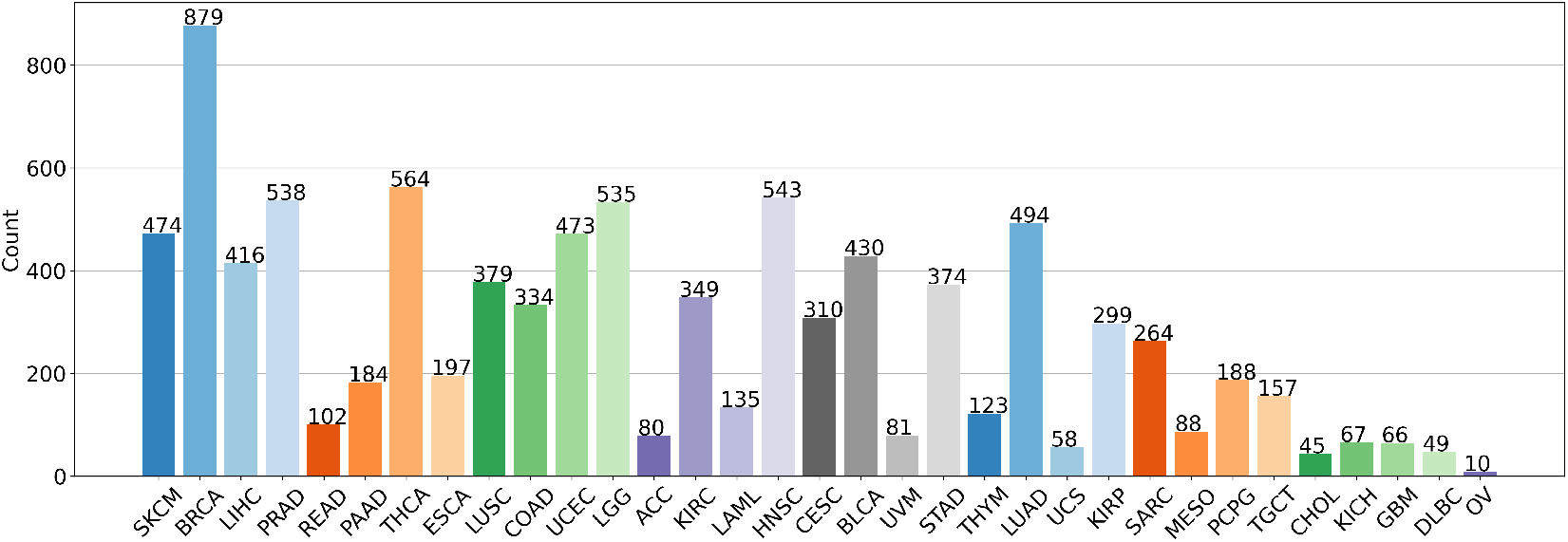
Pan-cancer classification label distribution.

### C.2. Downstream tasks benchmarks

In addition to Table 1 (cancer-type classification) and Table 3 (survival analysis), an exhaustive benchmark including other representation models for RNA-seq and DNA methylation was conducted:

- Multiple Factor Analysis (MFA) (Sánchez et al., 2012), using a latent space of dimension 256.
- Non-negative Matrix Factorization (NMF) (Lee & Seung, 1999), with the same latent space dimension as for MFA.
- *OmiEmbed* (Zhang et al., 2021b): a unified multi-task deep learning framework for multi-omics data based on Variational Auto-Encoders (Kingma & Welling, 2013) from early integrated omics.
- *IntegrAO* (Ma et al., 2024): an unsupervised framework based on Graph Neural Networks (Scarselli et al., 2008) for integrating incomplete multi-omics data, tailored for classification and survival task.

Multiple Factor Analysis and Non-negative Matrix Factorization features are then fed to a Support Vector Machine (SVM) for the cancer-type classification task and to a Cox proportional hazards model for the survival analysis task. The results are presented in Table 10 and Table 11. We also report in Table 12 the complete benchmark on the zero-shot classification and clustering tasks which includes *CustOmics* model.

**Table 10.** Full benchmark on cancer-type classification. **Bold**: best; <u>underline</u>: second-best.

| Model | Modality | Macro-F1 | Weighted-F1 |
| --- | --- | --- | --- |
| *Unimodal encoders* |  |  |  |
| BulkRNABert | RNA-seq | 0.918 ± 0.008 | 0.943 ± 0.004 |
| scGPT | RNA-seq | 0.793 ± 0.009 | 0.831 ± 0.008 |
| MethFormer | Methylation | 0.917 ± 0.008 | 0.931 ± 0.006 |
| *Classical multi-omics integration* |  |  |  |
| MFA | Bimodal | 0.753 ± 0.013 | 0.848 ± 0.008 |
| NMF | Bimodal | 0.725 ± 0.011 | 0.827 ± 0.006 |
| MOFA | Bimodal | 0.789 ± 0.012 | 0.852 ± 0.007 |
| *Deep learning multi-omics integration* |  |  |  |
| IntegrAO | Bimodal | 0.912 ± 0.005 | 0.911 ± 0.015 |
| OmiEmbed | Bimodal | 0.919 ± 0.004 | 0.922 ± 0.016 |
| CustOmics (probing) | Bimodal | 0.887 ± 0.065 | 0.911 ± 0.088 |
| CustOmics (end-to-end) | Bimodal | 0.922 ± 0.006 | <u>0.946 ± 0.006</u> |
| SeNMo | Bimodal | 0.900 ± 0.013 | 0.922 ± 0.014 |
| GAT | Bimodal | 0.882 ± 0.008 | 0.915 ± 0.004 |
| *Late integration (pre-trained encoders)* |  |  |  |
| MultiSurv | Bimodal | 0.911 ± 0.005 | 0.932 ± 0.005 |
| Late integration (concatenation) | Bimodal | 0.928 ± 0.008 | 0.945 ± 0.007 |
| Late integration (cross-attention) | Bimodal | <u>0.929 ± 0.005</u> | 0.945 ± 0.002 |
| *Joint bimodal pre-training (MOJO variants)* |  |  |  |
| MOJO-Transformer (no pre-training) | Bimodal | 0.802 ± 0.008 | 0.827 ± 0.010 |
| MOJO-Transformer (probing) | Bimodal | 0.850 ± 0.014 | 0.892 ± 0.009 |
| MOJO-Transformer | Bimodal | 0.925 ± 0.005 | 0.942 ± 0.003 |
| MOJO (no pre-training) | Bimodal | 0.835 ± 0.015 | 0.891 ± 0.006 |
| MOJO (probing) | Bimodal | 0.928 ± 0.009 | 0.945 ± 0.006 |
| MOJO | Bimodal | **0.935 ± 0.007** | **0.952 ± 0.006** |

**Table 11.** Full benchmark on pan-cancer survival analysis. **Bold**: best; <u>underline</u>: second-best.

| Model | Modality | C-index | Weighted C-index |
| --- | --- | --- | --- |
| *Unimodal encoders* |  |  |  |
| BulkRNABert | RNA-seq | $0.750 \pm 0.004$ | $0.657 \pm 0.011$ |
| scGPT | RNA-seq | $0.720 \pm 0.005$ | $0.604 \pm 0.020$ |
| MethFormer | Methylation | $0.735 \pm 0.006$ | $0.618 \pm 0.017$ |
| *Classical multi-omics integration* |  |  |  |
| MFA | Bimodal | $0.616 \pm 0.033$ | $0.593 \pm 0.016$ |
| NMF | Bimodal | $0.616 \pm 0.040$ | $0.591 \pm 0.025$ |
| MOFA | Bimodal | $0.648 \pm 0.037$ | $0.601 \pm 0.022$ |
| *Deep learning multi-omics integration* |  |  |  |
| IntegrAO | Bimodal | $0.710 \pm 0.008$ | $0.624 \pm 0.006$ |
| OmiEmbed | Bimodal | $0.736 \pm 0.006$ | $0.631 \pm 0.007$ |
| CustOmics | Bimodal | $0.686 \pm 0.018$ | $0.639 \pm 0.099$ |
| SeNMos | Bimodal | $0.745 \pm 0.004$ | <u>$0.640 \pm 0.017$</u> |
| GAT | Bimodal | <u>$0.757 \pm 0.007$</u> | $0.619 \pm 0.013$ |
| *Late integration (pre-trained encoders)* |  |  |  |
| Late integration (concat.) | Bimodal | $0.756 \pm 0.004$ | $0.653 \pm 0.011$ |
| MultiSurv | Bimodal | $0.742 \pm 0.002$ | $0.636 \pm 0.011$ |
| *Joint bimodal pre-training (MOJO variants)* |  |  |  |
| MOJO-Transformer | Bimodal | $0.757 \pm 0.006$ | $0.657 \pm 0.007$ |
| MOJO | Bimodal | **$0.771 \pm 0.006$** | **$0.670 \pm 0.009$** |

**Table 12.** Full benchmark on zero-shot classification and clustering results on pan-cancer and PAM50 tasks. (Acc. = Accuracy, NMI = Normalized Mutual Information, ARI = Adjusted Rand Index).

| Task | Metric | MOJO | Late integration | CustOmics |
| --- | --- | --- | --- | --- |
| PAM50 | Acc. | **0.777** | 0.763 | 0.765 |
|  | NMI | **0.345** | 0.291 | 0.311 |
|  | ARI | **0.213** | 0.154 | 0.176 |
| Pan-cancer | Acc. | **0.928** | 0.870 | 0.905 |
|  | NMI | **0.862** | 0.771 | 0.830 |
|  | ARI | **0.756** | 0.620 | 0.699 |

### C.3. *IA*^3^ fine-tuning for Survival Analysis

For classification, we apply *IA*^3^ fine-tuning to *MOJO, BulkRNABert*, and *MethFormer*. For survival analysis, we use probing only (frozen encoders) due to batch size requirements. To validate that this choice does not disadvantage MOJO, we compare probing vs. *IA*^3^ fine-tuning for survival in Table 13.

**Table 13.** Effect of *IA*^3^ fine-tuning on survival analysis. Differences are within the margin of error, confirming that probing is sufficient for this task.

| Model | $IA^3$ | C-index | Weighted C-index |
| --- | --- | --- | --- |
| MOJO | ✗ | $0.771 \pm 0.006$ | $0.670 \pm 0.009$ |
| MOJO | ✓ | $0.771 \pm 0.007$ | $0.670 \pm 0.010$ |
| BulkRNABert | ✗ | $0.749 \pm 0.003$ | $0.654 \pm 0.014$ |
| BulkRNABert | ✓ | $0.752 \pm 0.004$ | $0.657 \pm 0.010$ |
| MethFormer | ✗ | $0.736 \pm 0.006$ | $0.622 \pm 0.014$ |
| MethFormer | ✓ | $0.737 \pm 0.006$ | $0.620 \pm 0.016$ |

### C.4. Bimodal embeddings t-SNE visualisations

**Figure 10.**
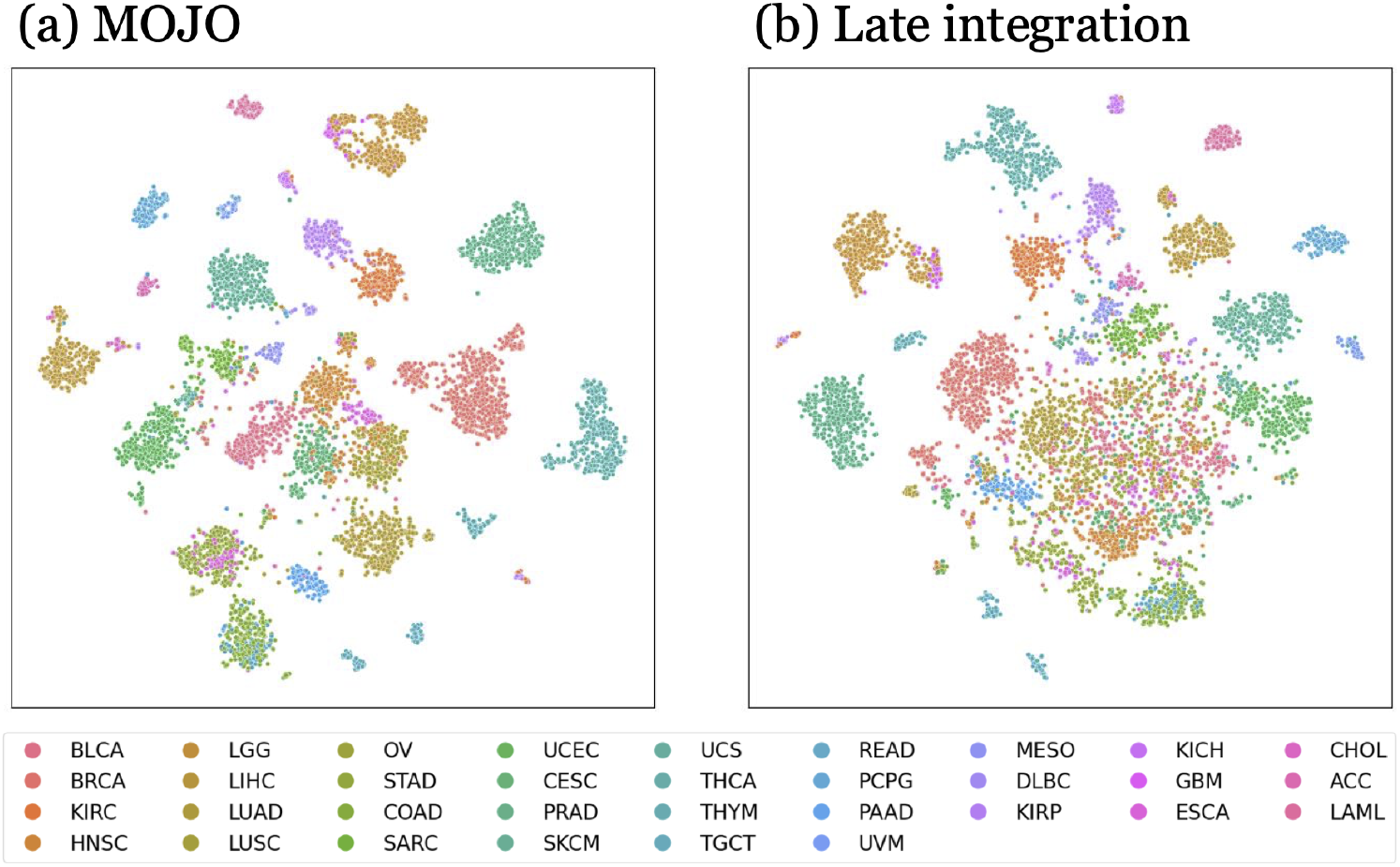
Pan-cancer version of the t-SNE representation of *MOJO* and *Late integration* embeddings, colored by cancer-type.

### C.5. Kaplan-Meier curves

**Figure 11.**
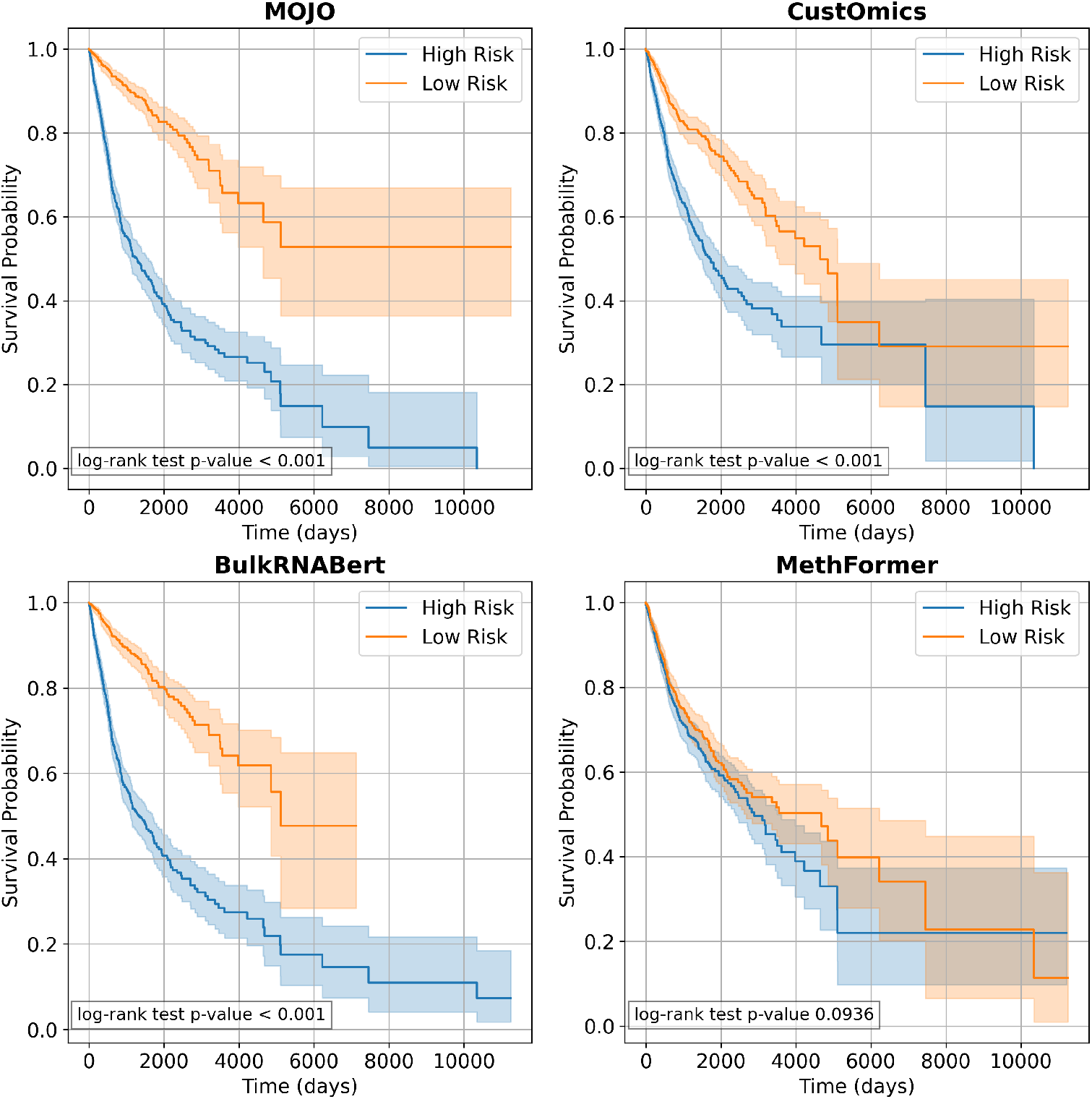
Kaplan-Meier curve for pan-cancer survival models for four models: *MOJO, CustOmics, BulkRNABert, MethFormer*. MOJO shows better patient stratification. “*Low Risk*” and “*High Risk*” are defined from the median of the log risk computed on the test set for each model.

## D. Region-Aware Methylation Aggregation

### D.1. Methodology

To map site-level methylation data to gene-level features compatible with bulk RNA-seq, in addition to the simple average of methylation values used throughout our work, we implemented a region-aware weighted aggregation strategy that accounts for the functional heterogeneity of genomic regions. We assigned varying weights to CpG sites based on their genomic location, reflecting the empirically observed distance-dependent impact of methylation on gene expression. Promoter-proximal regions (TSS200, TSS1500) received the highest weights (3.0, 2.5) due to their strong negative correlation with gene expression and established role in transcriptional silencing (Weber et al., 2007; Jones & Baylin, 2007), while 5’UTR and first exon sites received intermediate weights (2.0) based on their impact on transcriptional silencing and inhibition of elongation (Brenet et al., 2011). Gene body and 3’UTR regions served as baseline (weight = 1.0) given their weaker and often positive correlation with transcription (Ball et al., 2009), creating a biologically grounded weighting scheme that prioritizes functionally critical regulatory sites.

Additionally, we incorporated CpG island context weights to capture the differential regulatory potential of CpG sites based on their proximity to CpG islands. CpG islands (weight = 2.0) are strongly associated with gene regulation and promoter activity (Deaton & Bird, 2011), while CpG shores (weight = 1.5) have been shown to exhibit tissue-specific differential methylation with functional consequences (Irizarry et al., 2009). CpG shelves (weight = 1.0) and open sea regions (weight = 0.8) received lower weights reflecting their more variable and less consistent associations with gene expression.

The final weight for each CpG site *s* associated with gene *g* was computed as:

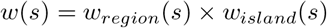

where *w*_*region*_(*s*) is the genomic region weight and *w*_*island*_(*s*) is the CpG island context weight, as detailed in Table 14.

**Table 14.** Weights assigned to CpG sites based on genomic region and CpG island context for region-aware methylation aggregation.

| Genomic Region | $w_{region}$ | CpG Island Context | $w_{island}$ |
| --- | --- | --- | --- |
| TSS200 | 3.0 | Island | 2.0 |
| TSS1500 | 2.5 | Shore | 1.5 |
| 5'UTR | 2.0 | Shelf | 1.0 |
| 1st Exon | 2.0 | Open Sea | 0.8 |
| Gene Body | 1.0 |  |  |
| 3'UTR | 1.0 |  |  |

The weighted methylation value for each gene was then computed as:

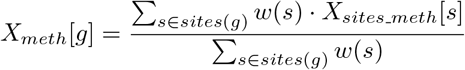

where *sites*(*g*) represents all CpG sites mapped to gene *g* according to the Illumina HumanMethylation450 BeadChip annotations (Bibikova et al., 2011; Sandoval et al., 2011), and *X*_*sites*_*_*_*meth*_[*s*] is the beta value for site *s*.

### D.2. Ablation study

We benchmark the weighted average of methylation value against the simple average on the pancancer cancer-type classification (Table 15) and survival analysis (Table 16) tasks.

**Table 15.** Ablation study on weighted average of methylation beta values for cancer type classification.

| Non-weighted average of methylation beta value |  | Weighted average of methylation beta value |  |
| --- | --- | --- | --- |
| Model | Test Weighted-F1 | Model | Test Weighted-F1 |
| BulkRNABert* | 0.943 ± 0.004 | BulkRNABert* | — |
| scGPT* | 0.831 ± 0.008 | scGPT* | — |
| MethFormer | 0.931 ± 0.006 | MethFormer | 0.935 ± 0.005 |
| MFA | 0.848 ± 0.008 | MFA | 0.850 ± 0.009 |
| NMF | 0.827 ± 0.006 | NMF | 0.824 ± 0.006 |
| MOFA | 0.852 ± 0.007 | MOFA | 0.849 ± 0.008 |
| IntegrAO | 0.911 ± 0.015 | IntegrAO | 0.944 ± 0.015 |
| OmiEmbed | 0.922 ± 0.016 | OmiEmbed | 0.925 ± 0.017 |
| Late integration (concat.) | 0.945 ± 0.007 | Late int. (concat.) | 0.940 ± 0.005 |
| Late integration (cross-att.) | 0.945 ± 0.002 | Late int. (cross-att.) | 0.941 ± 0.004 |
| Late integration (max.) | 0.911 ± 0.005 | Late int. (max.) | 0.930 ± 0.010 |
| CustOmics (probing) | 0.911 ± 0.088 | CustOmics (probing) | 0.893 ± 0.071 |
| CustOmics (e2e**) | 0.946 ± 0.006 | CustOmics (e2e**) | 0.941 ± 0.003 |
| SeNMo | 0.922 ± 0.014 | SeNMo | 0.937 ± 0.006 |
| GAT | 0.915 ± 0.004 | GAT | 0.920 ± 0.004 |
| MOJO-Transformer (probing) | 0.892 ± 0.009 | MOJO-Transformer (probing) | 0.896 ± 0.007 |
| MOJO-Transformer | 0.942 ± 0.003 | MOJO-Transformer | 0.938 ± 0.010 |
| MOJO (probing) | 0.945 ± 0.006 | MOJO (probing) | 0.950 ± 0.006 |
| MOJO | **0.952 ± 0.006** | MOJO | **0.951 ± 0.005** |

\*BulkRNABert and scGPT use only RNA data and are unaffected by methylation preprocessing. \*\*e2e = end-to-end

**Table 16.** Ablation study on weighted average of methylation beta values for survival analysis.

| Non-weighted average of methylation beta value |  | Weighted average of methylation beta value |  |
| --- | --- | --- | --- |
| Model | Weighted C-index | Model | Weighted C-index |
| BulkRNABert* | $0.657 \pm 0.011$ | BulkRNABert* | — |
| scGPT* | $0.604 \pm 0.020$ | scGPT* | — |
| MethFormer | $0.618 \pm 0.017$ | MethFormer | $0.622 \pm 0.018$ |
| MFA | $0.593 \pm 0.016$ | MFA | $0.594 \pm 0.018$ |
| NMF | $0.591 \pm 0.025$ | NMF | $0.590 \pm 0.026$ |
| MOFA | $0.601 \pm 0.022$ | MOFA | $0.598 \pm 0.024$ |
| IntegrAO | $0.624 \pm 0.006$ | IntegrAO | $0.625 \pm 0.006$ |
| OmiEmbed | $0.631 \pm 0.007$ | OmiEmbed | $0.633 \pm 0.007$ |
| CustOmics | $0.639 \pm 0.099$ | CustOmics | $0.642 \pm 0.023$ |
| Late integration (concat.) | $0.653 \pm 0.011$ | Late integration (concat.) | $0.652 \pm 0.016$ |
| MultiSurv | $0.636 \pm 0.011$ | MultiSurv | $0.635 \pm 0.021$ |
| SeNMo | $0.640 \pm 0.017$ | SeNMo | $0.644 \pm 0.021$ |
| GAT | $0.619 \pm 0.013$ | GAT | $0.615 \pm 0.018$ |
| MOJO-Transformer | $0.657 \pm 0.007$ | MOJO-Transformer | $0.641 \pm 0.010$ |
| MOJO | **$0.670 \pm 0.009$** | MOJO | **$0.654 \pm 0.018$** |

\*BulkRNABert and scGPT use only RNA data and are unaffected by methylation preprocessing.

While the region-aware weighted methylation aggregation strategy was designed to better capture the differential regulatory potential of CpG sites, our empirical evaluation reveals nuanced results. As shown in Tables 15 and 16, this biologically motivated weighting scheme does not yield improvements for the MOJO model in either cancer-type classification or survival prediction tasks; therefore, we retain the simpler average-based aggregation in our final model. This confirms that *MOJO* learns the relevant regulatory structure directly from the data, obviating the need for hand-crafted methylation priors. However, when examining the *MethFormer* baseline, we observe marginal improvements with region-aware aggregation. Furthermore, during pre-training, the region-aware scheme achieves consistently higher reconstruction accuracy compared to simple averaging (Figure 12). This improved reconstruction performance suggests that weighted aggregation, by accounting for the differential regulatory roles of promoters, gene bodies, and CpG islands, yields a more biologically grounded representation of methylation patterns. Nevertheless, these improvements do not transfer to the multimodal setting, indicating that the additional biological signal captured by region-aware weighting may be redundant with information already encoded within the RNA-seq modality. This finding highlights a broader challenge in multimodal learning: biologically motivated features that improve unimodal representation quality do not necessarily enhance multimodal fusion when modalities contain overlapping biological signals.

**Figure 12.**
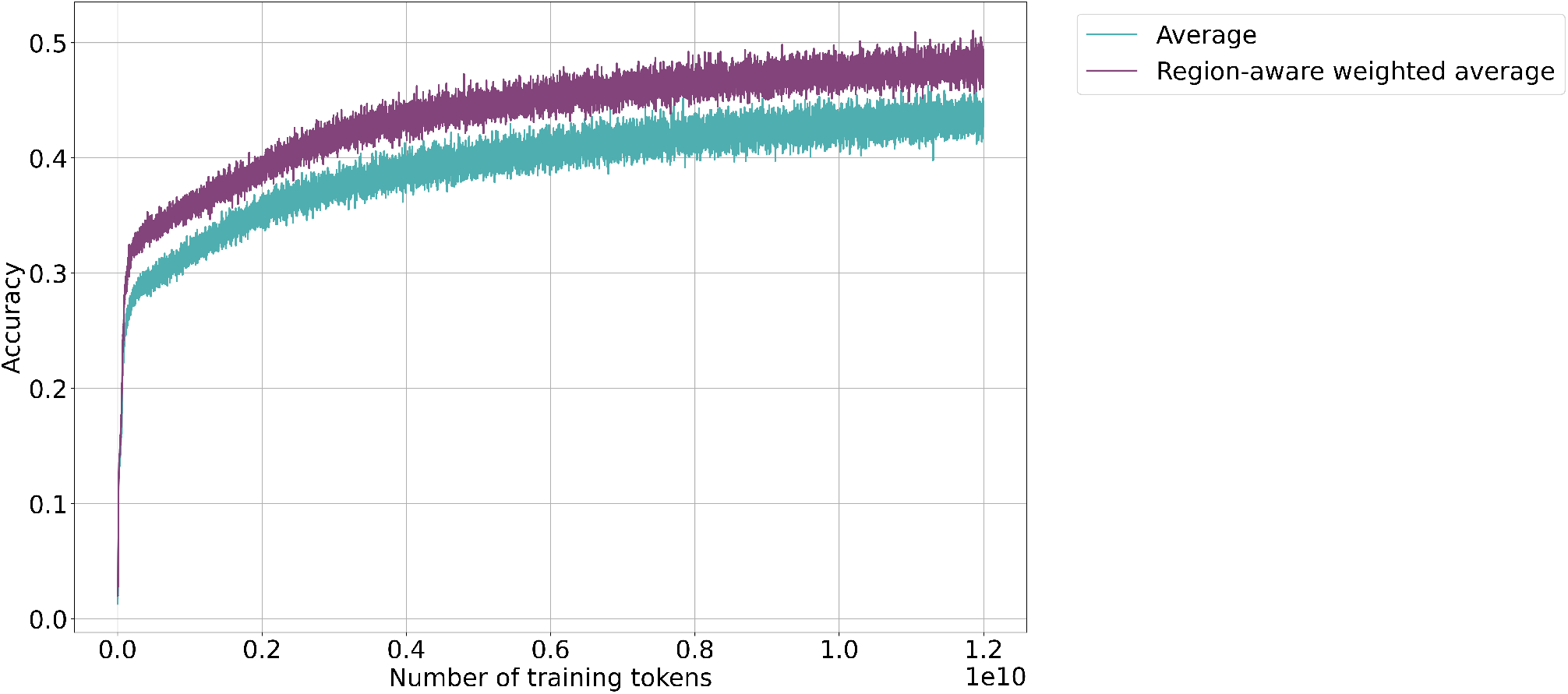
Comparison of *MethFormer* pre-training learning curves for the different methylation preprocessing schemes.

## E. Validation on external datasets

We evaluate the generalization of *MOJO* on two cohorts entirely held out from pre-training: the pediatric TARGET dataset and the ICGC cohort. Both datasets were preprocessed using identical pipelines to TCGA and evaluated on the same downstream tasks.

### E.2. TARGET

The TARGET (Therapeutically Applicable Research to Generate Effective Treatments, https://www.cancer.gov/ccg/research/genome-sequencing/target) dataset comprises primary solid tumor samples from the GDC portal with matched bulk RNA-seq and DNA methylation (Illumina 450k) data and survival information, resulting in 351 patients across four pediatric cancer types.

#### E.1.1. Dataset composition

**Table 17.** Composition of the TARGET dataset.

| Cohort | Cancer type | # Samples | # w/ survival | # Events | # Censored | # Event rate |
| --- | --- | --- | --- | --- | --- | --- |
| TARGET-CCSK | Clear Cell Sarcoma of the Kidney | 11 | 11 | 3 | 8 | 27.3% |
| TARGET-NBL | Neuroblastoma | 133 | 133 | 66 | 67 | 49.6% |
| TARGET-OS | Osteosarcoma | 85 | 82 | 29 | 53 | 35.4% |
| TARGET-WT | Wilms Tumor (Kidney) | 122 | 122 | 50 | 72 | 41.0% |
| **Total** | **All cohorts** | **351** | **348** | **148** | **200** | **42.5%** |

#### E.1.2. Batch effect analysis

We quantified cross-platform batch effects using three complementary metrics: (1) Silhouette scores to measure global dataset separation (range: −1 to 1, with 0 indicating complete mixing), (2) the *k*-nearest neighbor Batch Effect Test (kBET) to assess whether batch labels are randomly distributed within local neighborhoods (range: 0 to 1, with 1 indicating perfect mixing), and (3) the Pearson correlation of mean bulk RNA-seq and DNA methylation levels between platforms. Our analysis revealed weak batch effects (bulk RNA-seq: Silhouette= 0.19, kBET= 0.95, *r* = 0.86; DNA methylation: Silhouette= 0.12, kBET= 0.92, *r* = 0.96). PCA visualization further showed substantial overlap between TCGA and TARGET samples (Figure 13). The high kBET scores and strong correlations demonstrate excellent local mixing and overall concordance, validating cross-platform generalization.

**Figure 13.**
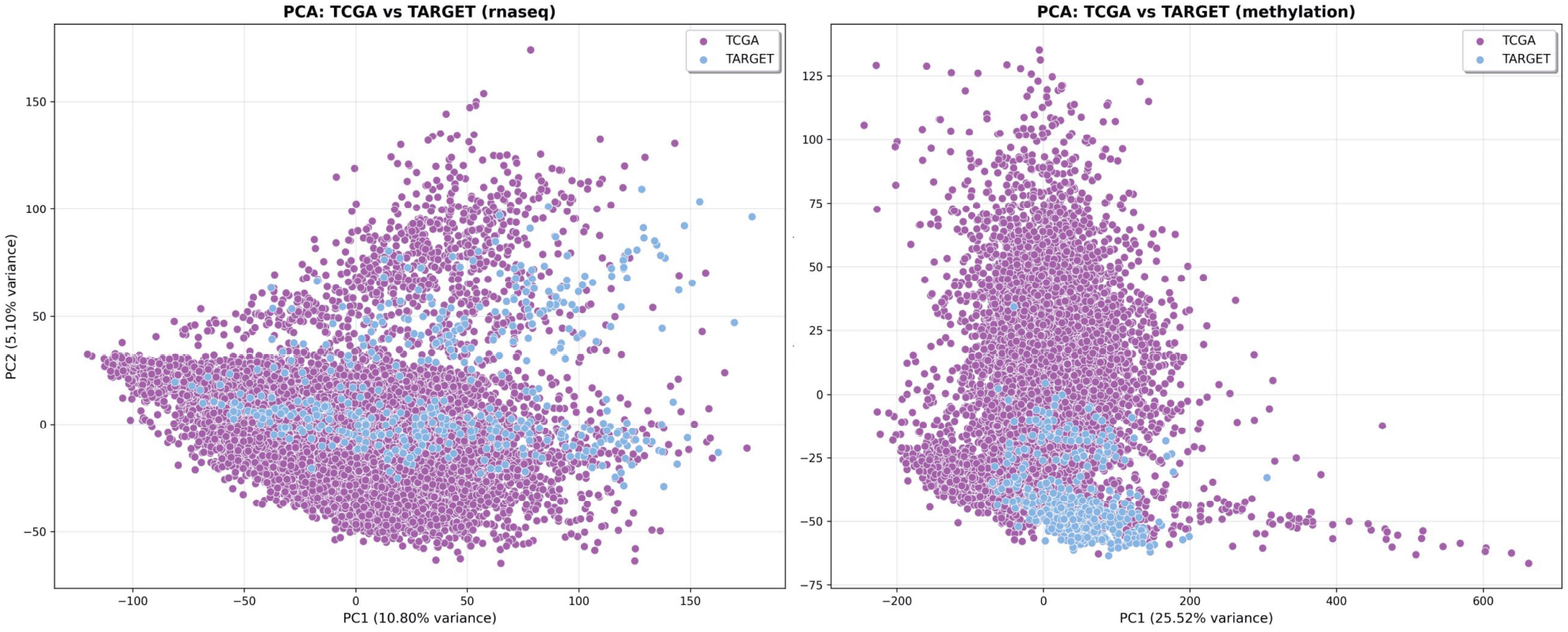
Bulk RNA-seq and DNA methylation PCA plots — comparison of TCGA and TARGET datasets.

#### E.1.3. Results

We evaluated models on two tasks: survival analysis (348 patients with follow-up) and zero-shot classification and clustering, discriminating the four cancer types without any fine-tuning on TARGET.

**Table 18.** External validation on the TARGET dataset.

| (a) Survival analysis |  |  | (b) Zero-shot classification and clustering |  |  |
| --- | --- | --- | --- | --- | --- |
| Model | C-index | Weighted C-index | Metric | MOJO | Late integration |
| BulkRNABert | **0.601 ± 0.042** | 0.599 ± 0.039 | Acc. | **0.992** | 0.991 |
| MethFormer | 0.548 ± 0.027 | 0.540 ± 0.042 | NMI | **0.949** | 0.794 |
| MOJO | **0.601 ± 0.023** | **0.606 ± 0.042** | ARI | **0.946** | 0.706 |

### E.2. ICGC External Validation Cohort

The ICGC (International Cancer Genome Consortium, https://dcc.icgc.org) cohort was assembled from all non-US ICGC projects with publicly available paired bulk RNA-seq and DNA methylation (Illumina 450k) data, after excluding any donors overlapping with TCGA.

#### E.2.1. Dataset composition

**Table 19.**
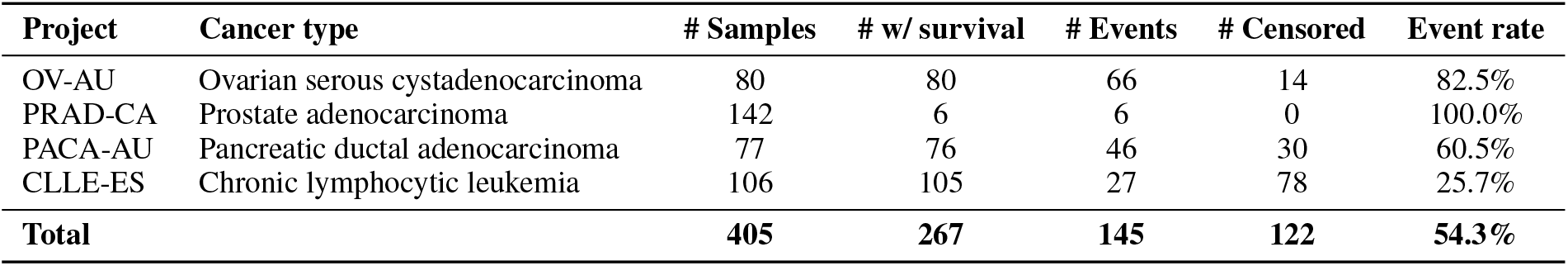
Composition of the ICGC external validation cohort. Event rate is the proportion deceased among samples with survival information.

| Project | Cancer type | # Samples | # w/ survival | # Events | # Censored | Event rate |
| --- | --- | --- | --- | --- | --- | --- |
| OV-AU | Ovarian serous cystadenocarcinoma | 80 | 80 | 66 | 14 | 82.5% |
| PRAD-CA | Prostate adenocarcinoma | 142 | 6 | 6 | 0 | 100.0% |
| PACA-AU | Pancreatic ductal adenocarcinoma | 77 | 76 | 46 | 30 | 60.5% |
| CLLE-ES | Chronic lymphocytic leukemia | 106 | 105 | 27 | 78 | 25.7% |
| **Total** |  | **405** | **267** | **145** | **122** | **54.3%** |

For survival analysis, PRAD-CA was excluded: with only 6 of 142 donors having usable follow-up in ICGC (all 6 deceased), the cohort provides insufficient signal for reliable survival modelling. The remaining three cohorts yield 261 donors with a 53% event rate and a median follow-up of 1,162 days.

#### E.2.2. Results

We evaluated models on survival analysis (261 donors from OV-AU, PACA-AU, and CLLE-ES), and cancer-type classification (distinguishing the four ICGC lineages across all 405 bimodal samples). Classification performance was uninformative in this setting: all three models trivially reached a macro-*F*_1_ of 1.000, as these four cancer lineages are biologically distant and easily separable by any omics model. The survival results (Table 20) confirm that MOJO’s performance advantage over unimodal baselines generalizes to fully out-of-domain data.

**Table 20.** Survival analysis on the ICGC external cohort (261 donors; 3 cohorts: OV-AU, PACA-AU, CLLE-ES; 54% event rate). **Bold**: best; <u>underline</u>: second-best.

| Model | C-index | Weighted C-index |
| --- | --- | --- |
| BulkRNABert | 0.805 ± 0.036 | 0.610 ± 0.087 |
| MethFormer | <u>0.810 ± 0.027</u> | <u>0.628 ± 0.062</u> |
| MOJO | **0.823 ± 0.019** | **0.643 ± 0.032** |

## F. Interpretability results

In this section, we provide additional results for the interpretability section 5.4. Figure 14 provides a swarm plot of the SHAP values for the *Luminal B* subtype of BRCA. The results demonstrate a strong monotonic impact of the most influential dimensions on *Luminal B* subtype predictions. For a given dimension, its high values are associated with SHAP values of a consistent sign, while its low values are associated with the opposite, indicating that the model has learned distinct predictive patterns. Figure 15 provides the pre-rank GSEA results for all four BRCA subtypes. We only kept the enriched pathways with a false detection rate (FDR) q-value less than 0.25 (Subramanian et al., 2005) and we present only pathways with positive enrichment scores.

**Figure 14.**
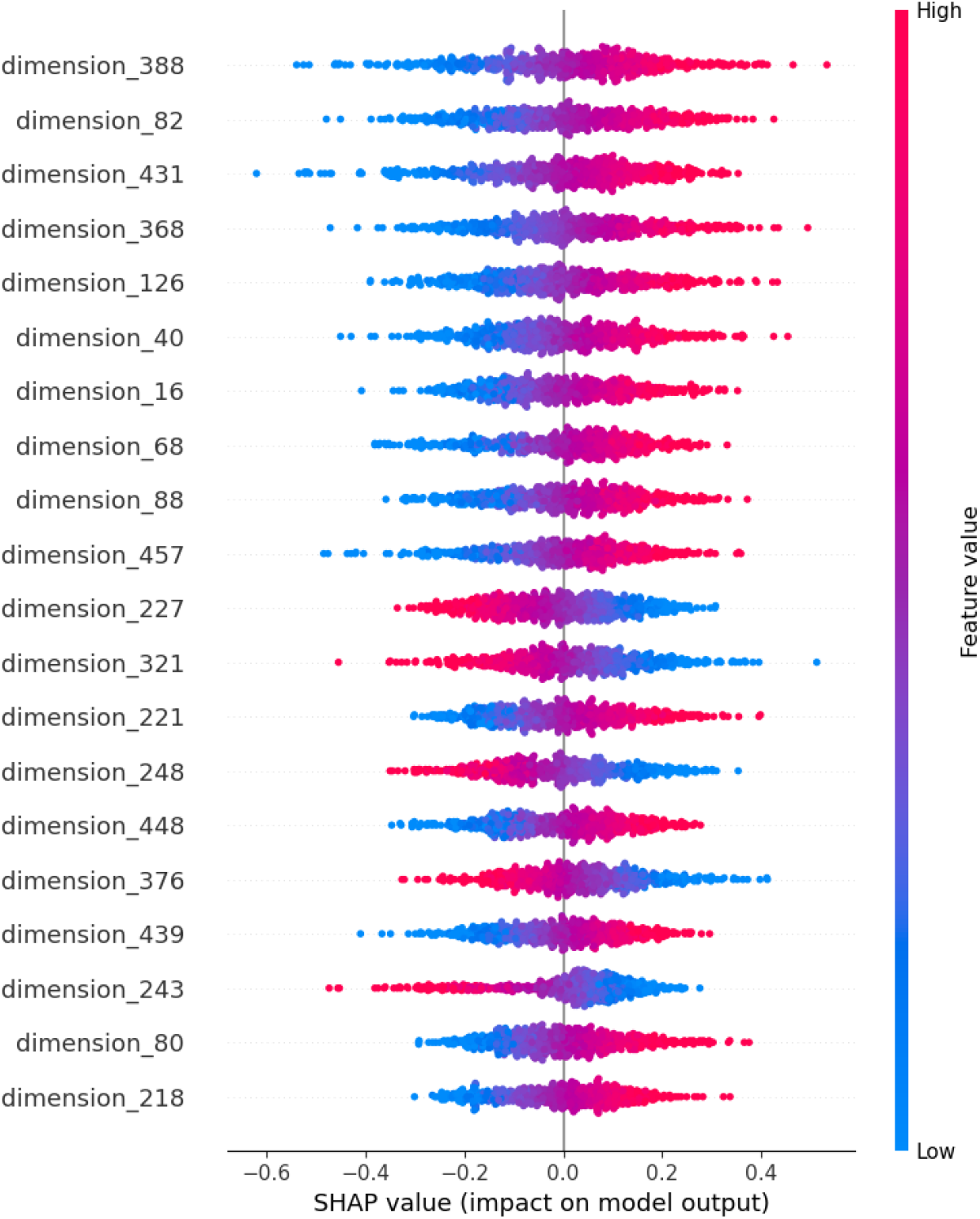
Shapley Values swarmplot for *LumB* sub-type of BRCA cancer.

**Figure 15.**
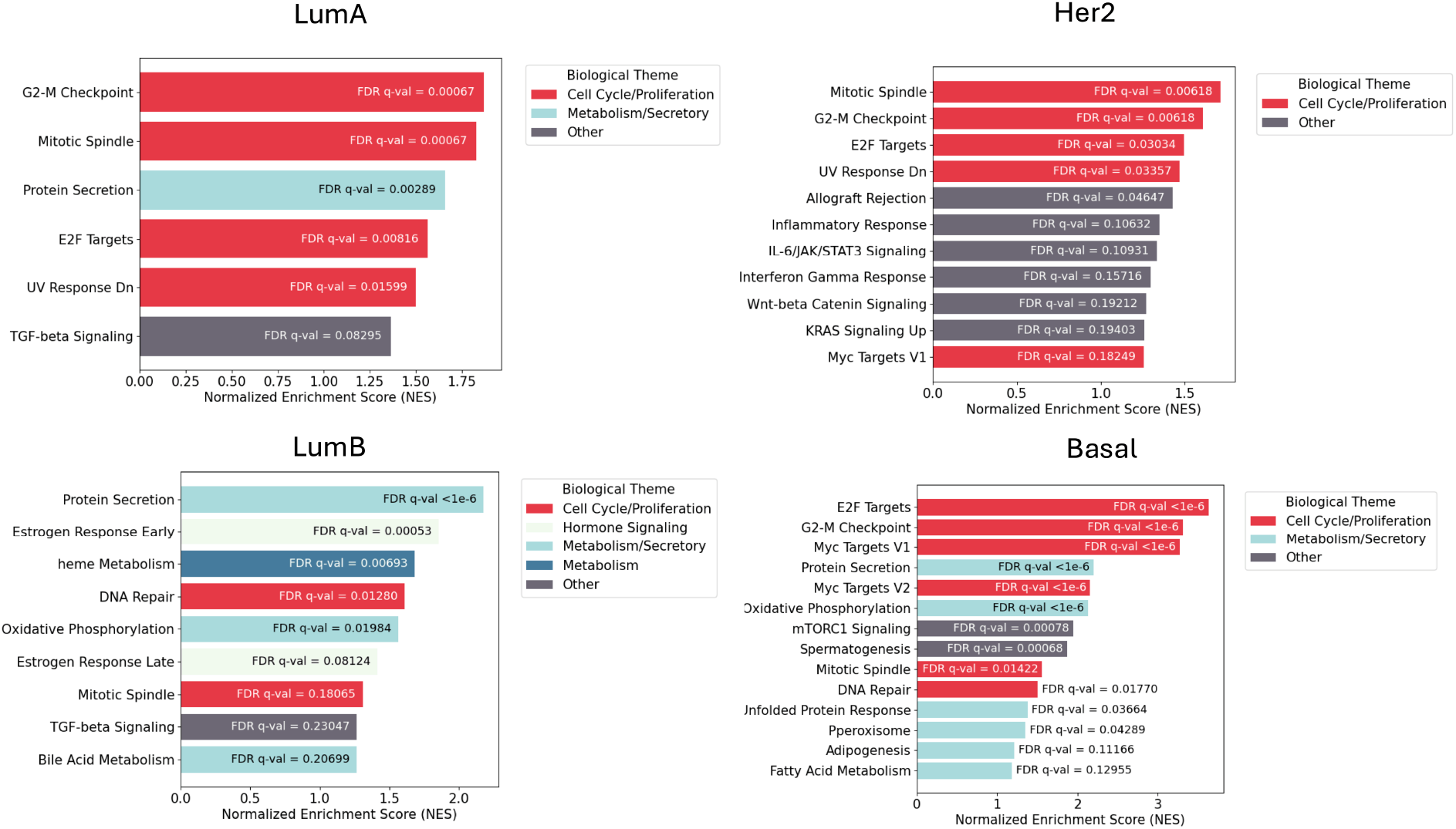
Pre-rank gene set enrichment analysis results for all subtypes of BRCA from bulk RNA-seq correlations with most predictive *MOJO*’s embedding dimensions.

## G. Cross-Platform Transferability to 27k Arrays

The Illumina HumanMethylation27k BeadChip is a legacy platform still prevalent in older public cohorts. Unlike the 450k array used to pre-train *MOJO*, 27k arrays cover only ~ 27,000 CpG sites with a strong promoter bias, resulting in systematically different gene-level beta-values. Most methylation models trained on 450k data are therefore incompatible with 27k data without retraining. MOJO’s gene-level aggregation makes it inherently array-agnostic: switching platforms simply changes which probes contribute to each gene’s mean, without requiring any architectural change. We evaluate this property here and provide the full results supporting Section 6.1.

### Dataset construction

We generated an in-silico 27k dataset by filtering our TCGA 450k data to the 27k microarray probe IDs, maintaining exact patients for a 1-to-1 comparison. Gene-level features are computed using the same simple probe averaging as for 450k data. Of the 25,887 27k probes mapping to MOJO’s gene vocabulary, 13,726/17,116 genes (80.2%) obtain at least one probe; genes without coverage receive the <MASK> token.

### Distribution shift quantification

The 27k array’s strong promoter bias causes a systematic mean shift of −0.115 in gene-level beta-values relative to 450k averages, inducing a severe distribution shift that degrades zero-shot model performance.

### Evaluation strategies

We compare three adaptation strategies, all with entirely frozen model backbones except for the last:

- *Zero-shot*: the 27k input is fed directly to the model without any adaptation.
- *Rescaled*: for each train/test split, a per-gene z-score normalization is applied using the mean and standard deviation of each covered gene computed over the 450k training fold, re-scaling values to the 450k range. No retraining is required.
- *Head fine-tuning*: task heads are retrained on 27k data while the backbone remains frozen.

**Table 21.** 27k array transferability under different adaptation strategies for CustOmics and MOJO. **Bold**: best per strategy and metric.

| Evaluation setting | Model | Cancer-type Classification |  | Survival Analysis |  |
| --- | --- | --- | --- | --- | --- |
|  |  | Macro-F1 | Weighted-F1 | C-index | Weighted C-index |
| 450k (Baseline) | CustOmics | $0.922 \pm 0.006$ | $0.946 \pm 0.006$ | $0.686 \pm 0.018$ | $0.639 \pm 0.099$ |
|  | *MOJO* | **$0.935 \pm 0.007$** | **$0.952 \pm 0.006$** | **$0.771 \pm 0.006$** | **$0.670 \pm 0.009$** |
| 27k (Zero-shot) | CustOmics | $0.226 \pm 0.001$ | $0.336 \pm 0.002$ | $0.509 \pm 0.008$ | $0.515 \pm 0.006$ |
|  | *MOJO* | **$0.734 \pm 0.024$** | **$0.824 \pm 0.015$** | **$0.717 \pm 0.018$** | **$0.639 \pm 0.025$** |
| 27k (Rescaled) | CustOmics | $0.761 \pm 0.012$ | $0.818 \pm 0.007$ | $0.633 \pm 0.051$ | $0.612 \pm 0.044$ |
|  | *MOJO* | **$0.909 \pm 0.007$** | **$0.940 \pm 0.003$** | **$0.765 \pm 0.007$** | **$0.664 \pm 0.016$** |
| 27k (Head fine-tuning) | CustOmics | $0.790 \pm 0.010$ | $0.837 \pm 0.004$ | $0.657 \pm 0.054$ | $0.629 \pm 0.028$ |
|  | *MOJO* | **$0.913 \pm 0.012$** | **$0.941 \pm 0.005$** | **$0.768 \pm 0.008$** | **$0.660 \pm 0.022$** |

The results highlight *MOJO*’s robustness at every level of adaptation. Under zero-shot evaluation, *CustOmics* collapses (macro-*F*_1_: 0.226) while MOJO retains strong performance (0.734), even outperforming fine-tuned *CustOmics* on survival (C-index 0.717 vs. 0.657). Z-score rescaling alone recovers 97% of *MOJO*’s 450k macro-*F*_1_ and 99% of its C-index, with head fine-tuning providing only marginal additional gains. While rescaling may be sensitive to batch effects in real-world deployment, making head fine-tuning the safer practical choice, both strategies confirm that *MOJO* captures highly transferable representations, making legacy 27k cohorts directly usable without re-pre-training or architectural changes.

## H. Missing modalities experiments

### H.1. Mutual Information computation

Algorithm 1 details the procedure for computing the mutual information loss for a single example. Figure 16 provides the complete results from the missing modalities experiments (when either RNA-seq or methylation is missing), while Table 22 provides exact figures.

**Figure 16.**
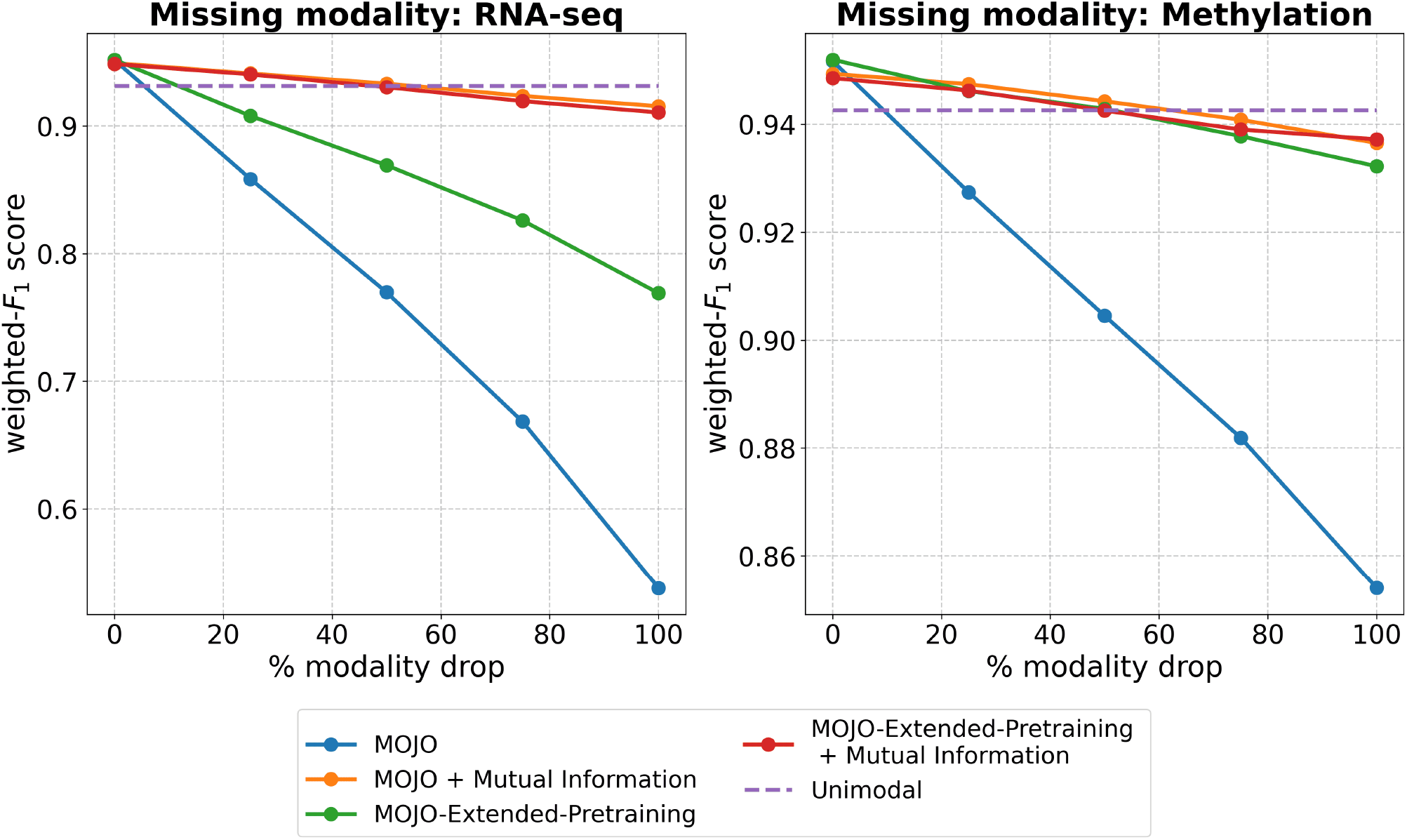
Missing modalities experimental results. Test weighted-*F*_1_ score for the pan-cancer classification is reported for different methods to handle the absence of a modality in x% of the samples (left: RNA-seq, right: Methylation). Unimodal models are respectively *MethFormer* and *BulkRNABert* when RNA-seq or Methylation is missing.

#### Algorithm 1 Mutual information auxiliary (MI) loss

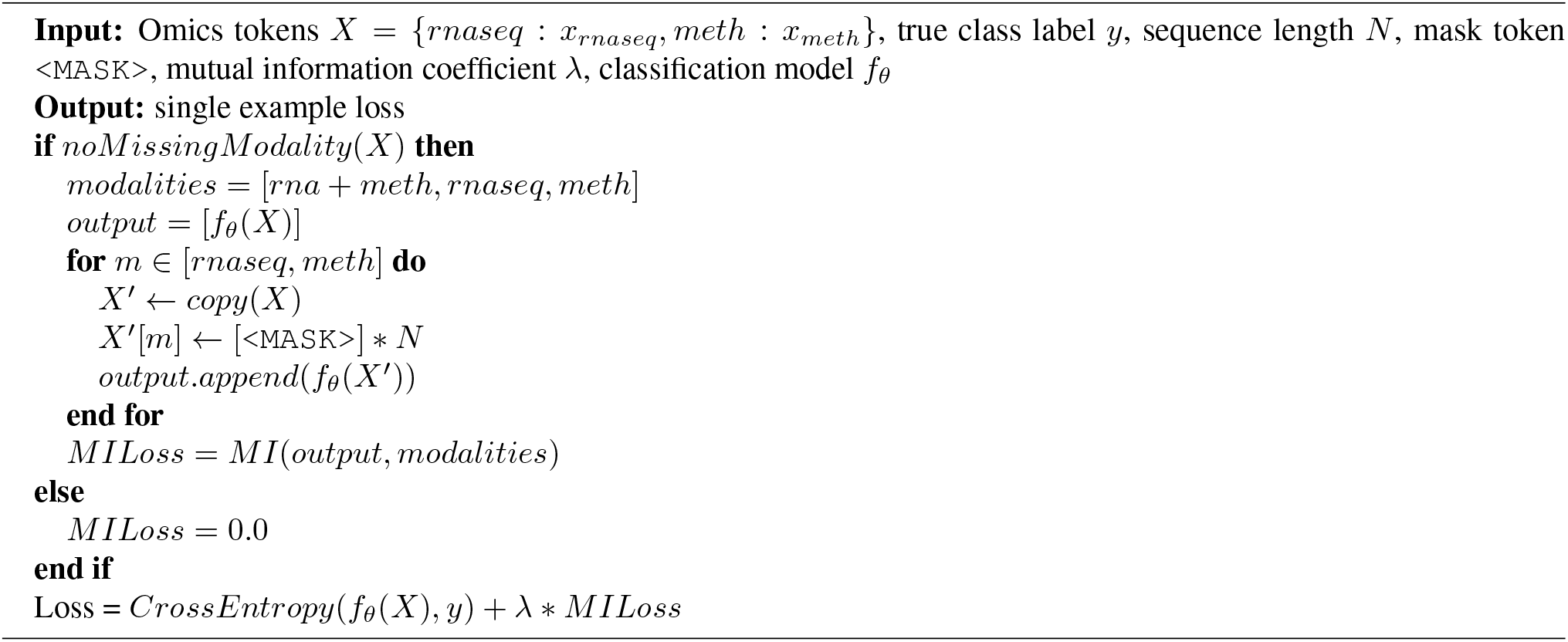

### H.2. Experimental results on mutual information

**Table 22.** Missing modalities experiment: cancer type classification.

| Model | Add mutual information | Drop modality (test time) | macro-F1 | test weighted-F1 |
| --- | --- | --- | --- | --- |
| BulkRNABert | ✗ | - | $0.918 \pm 0.008$ | $0.943 \pm 0.004$ |
| MethFormer | ✗ | - | $0.917 \pm 0.008$ | $0.931 \pm 0.006$ |
| MOJO | ✗ | - | $0.935 \pm 0.007$ | $0.952 \pm 0.006$ |
| MOJO | ✗ | Drop 100% of RNA-seq | $0.422 \pm 0.022$ | $0.538 \pm 0.025$ |
| MOJO | ✗ | Drop 100% of Methylation | $0.764 \pm 0.024$ | $0.854 \pm 0.011$ |
| MOJO | ✓ | - | $0.930 \pm 0.007$ | $0.949 \pm 0.004$ |
| MOJO | ✓ | Drop 100% of RNA-seq | $0.895 \pm 0.008$ | $0.916 \pm 0.007$ |
| MOJO | ✓ | Drop 100% of Methylation | $0.911 \pm 0.012$ | $0.937 \pm 0.008$ |
| MOJO-Extended-Pretraining | ✗ | - | $0.933 \pm 0.006$ | $0.952 \pm 0.003$ |
| MOJO-Extended-Pretraining | ✗ | Drop 100% of RNA-seq | $0.653 \pm 0.013$ | $0.769 \pm 0.004$ |
| MOJO-Extended-Pretraining | ✗ | Drop 100% of Methylation | $0.903 \pm 0.010$ | $0.932 \pm 0.005$ |
| MOJO-Extended-Pretraining | ✓ | - | $0.929 \pm 0.006$ | $0.949 \pm 0.005$ |
| MOJO-Extended-Pretraining | ✓ | Drop 100% of RNA-seq | $0.883 \pm 0.005$ | $0.911 \pm 0.004$ |
| MOJO-Extended-Pretraining | ✓ | Drop 100% of Methylation | $0.911 \pm 0.010$ | $0.937 \pm 0.006$ |

### H.3. Sensitivity analysis: Mutual Information coefficient *λ*

The missing-modality fine-tuning loss combines a primary cross-entropy term with an auxiliary mutual information term weighted by *λ* (see Section 7.2). We sweep *λ* ∈{1, 5, 10, 20, 50} to assess sensitivity to this hyperparameter, evaluating pan-cancer classification weighted-*F*_1_ under full bimodal input and under complete drop of each modality.

**Table 23.** Sensitivity of the mutual information auxiliary loss to *λ* ∈ {1, 5, 10, 20, 50}. Weighted-*F*_1_ on pan-cancer classification under no modality drop, 100% RNA-seq dropped, and 100% methylation dropped.

| $\lambda$ | Weighted-F1<br>No modality drop | Weighted-F1<br>Drop 100% RNA-seq | Weighted-F1<br>Drop 100% methylation |
| --- | --- | --- | --- |
| 1 | $0.953 \pm 0.006$ | $0.915 \pm 0.009$ | $0.939 \pm 0.006$ |
| 5 | $0.952 \pm 0.005$ | $0.918 \pm 0.008$ | $0.938 \pm 0.007$ |
| 10 | $0.949 \pm 0.004$ | $0.916 \pm 0.007$ | $0.937 \pm 0.008$ |
| 20 | $0.937 \pm 0.006$ | $0.908 \pm 0.007$ | $0.924 \pm 0.006$ |
| 50 | $0.912 \pm 0.004$ | $0.889 \pm 0.005$ | $0.900 \pm 0.006$ |

Performance is stable for *λ* ≤ 10, confirming our empirical choice. Values *λ* ≥ 20 degrade performance as the MI loss progressively dominates the primary cross-entropy signal.

### H.4. *MOJO* vs. *MOJO-Extended-Pretraining*: full-modality comparison

Figure 4 shows that *MOJO-Extended-Pretraining* improves unimodal fine-tuning (OV subtyping with RNA-seq only). To justify retaining standard *MOJO* as the primary model for bimodal tasks, we benchmark both models when both modalities are available in Table 24:

**Table 24.** MOJO vs. MOJO-Extended-Pretraining on full-modality downstream tasks. Results are statistically indistinguishable.

| Model | Cancer-type Classification |  | Survival Analysis |  |
| --- | --- | --- | --- | --- |
|  | Macro-F1 | Weighted-F1 | C-index | Weighted C-index |
| *MOJO* | $0.935 \pm 0.007$ | $0.952 \pm 0.006$ | $0.771 \pm 0.006$ | $0.670 \pm 0.009$ |
| *MOJO-Extended-Pretraining* | $0.933 \pm 0.006$ | $0.952 \pm 0.003$ | $0.768 \pm 0.008$ | $0.666 \pm 0.018$ |

Adding unimodal samples during pre-training provides no advantage for bimodal downstream tasks, while it specifically benefits scenarios where one modality is systematically absent during fine-tuning (as in OV sub-typing). We therefore retain standard *MOJO* for all bimodal benchmarks.

## References

Argelaguet, R., Velten, B., Arnol, D., Dietrich, S., Zenz, T., Marioni, J. C., Buettner, F., Huber, W., and Stegle, O. Multi-omics factor analysis—a framework for unsu-pervised integration of multi-omics data sets. Molecular systems biology, 14(6):e8124, 2018.

Avsec, Ž., Agarwal, V., Visentin, D., Ledsam, J. R., Grabska-Barwinska, A., Taylor, K. R., Assael, Y., Jumper, J., Kohli, P., and Kelley, D. R. Effective gene expression prediction from sequence by integrating long-range interactions. Nature methods, 18(10):1196–1203, 2021.

Ba, J. L., Kiros, J. R., and Hinton, G. E. Layer normalization. arXiv preprint arXiv:1607.06450, 2016.

Ball, M. P., Li, J. B., Gao, Y., Lee, J.-H., LeProust, E. M.,Park, I.-H., Xie, B., Daley, G. Q., and Church, G. M. Targeted and genome-scale strategies reveal gene-body methylation signatures in human cells. Nature Biotechnology, 27(4):361–368, 2009.

Benkirane, H., Pradat, Y., Michiels, S., and Cournede, P.-H. Customics: A versatile deep-learning based strategy for multi-omics integration. PLOS Computational Biology, 19(3):e1010921, 2023.

Bibikova, M., Barnes, B., Tsan, C., Ho, V., Klotzle, B., Le, J. M., Delano, D., Zhang, L., Schroth, G. P., Gunderson, K. L., et al. High density dna methylation array with single cpg site resolution. Genomics, 98(4):288–295, 2011.

Brenet, F., Moh, M., Funk, P., Feierstein, E., Viale, A. J., Socci, N. D., and Scandura, J. M. Dna methylation of the first exon is tightly linked to transcriptional silencing. PLoS ONE, 6(1):e14524, 2011.

Brixi, G., Durrant, M. G., Ku, J., Poli, M., Brockman, G., Chang, D., Gonzalez, G. A., King, S. H., Li, D. B., Merchant, A. T., et al. Genome modeling and design across all domains of life with evo 2. bioRxiv, pp. 2025–02, 2025.

Chen, Q., Zhang, J., Meng, R., Zhou, L., Li, Z., Feng, Q., and Shen, D. Modality-specific information disentanglement from multi-parametric mri for breast tumor segmentation and computer-aided diagnosis. IEEE Transactions on Medical Imaging, 43(5):1958–1971, 2024.

Chen, R. J., Lu, M. Y., Wang, J., Williamson, D. F., Rodig, S. J., Lindeman, N. I., and Mahmood, F. Pathomic fusion: an integrated framework for fusing histopathology and genomic features for cancer diagnosis and prognosis. IEEE Transactions on Medical Imaging, 41(4):757–770, 2020.

Ching, T., Zhu, X., and Garmire, L. X. Cox-nnet: an artificial neural network method for prognosis prediction of high-throughput omics data. PLoS computational biology, 14(4):e1006076, 2018.

Cortes, C. and Vapnik, V. Support-vector networks. Machine learning, 20:273–297, 1995.

Cox, D. R. Regression models and life-tables. Journal of the Royal Statistical Society: Series B (Methodological), 34(2):187–202, 1972.

Cui, H., Wang, C., Maan, H., Pang, K., Luo, F., Duan, N., and Wang, B. scgpt: toward building a foundation model for single-cell multi-omics using generative ai. Nature Methods, 21(8):1470–1480, 2024.

Dai, X. and Shen, L. Advances and trends in omics technology development. Frontiers in Medicine, 9:911861, 2022.

Dalla-Torre, H., Gonzalez, L., Mendoza-Revilla, J., Lopez Carranza, N., Grzywaczewski, A. H., Oteri, F., Dallago, C., Trop, E., de Almeida, B. P., Sirelkhatim, H., et al. Nucleotide transformer: building and evaluating robust foundation models for human genomics. Nature Methods, 22(2):287–297, 2025.

Deaton, A. M. and Bird, A. Cpg islands and the regulation of transcription. Genes & Development, 25(10):1010–1022, 2011.

Devlin, J., Chang, M.-W., Lee, K., and Toutanova, K. Bert: Pre-training of deep bidirectional transformers for language understanding. In Proceedings of the 2019 conference of the North American chapter of the association for computational linguistics: human language technologies, volume 1 (long and short papers), pp. 4171–4186, 2019.

Fertig, E. J., Lee, E., Pandey, N. B., and Popel, A. S. Analysis of gene expression of secreted factors associated with breast cancer metastases in breast cancer subtypes. Scientific reports, 5(1):12133, 2015.

Garau-Luis, J. J., Bordes, P., Gonzalez, L., Roller, M., de Almeida, B., Blum, C., Hexemer, L., Laurent, S., Lang, M., Pierrot, T., et al. Multi-modal transfer learning between biological foundation models. Advances in Neural Information Processing Systems, 37:78431–78450, 2024.

Gélard, M., Richard, G., Pierrot, T., and Cournede, P.-H. Bulkrnabert: Cancer prognosis from bulk rna-seq based language models. In Proceedings of the 4th Machine Learning for Health Symposium, volume 259 of Proceedings of Machine Learning Research, pp. 384–400. PMLR, 15–16 Dec 2025.

Gross, B., Dauvin, A., Cabeli, V., Kmetzsch, V., El Khoury, J., Dissez, G., Ouardini, K., Grouard, S., Davi, A., Loeb, R., et al. Robust evaluation of deep learning-based representation methods for survival and gene essentiality prediction on bulk rna-seq data. Scientific Reports, 14(1): 17064, 2024.

Harrell, F. E., Califf, R. M., Pryor, D. B., Lee, K. L., and Rosati, R. A. Evaluating the yield of medical tests. Jama, 247(18):2543–2546, 1982.

Ho, W. J., Erbe, R., Danilova, L., Phyo, Z., Bigelow, E., Stein-O’Brien, G., Thomas, D. L., Charmsaz, S., Gross, N., Woolman, S., et al. Multi-omic profiling of lung and liver tumor microenvironments of metastatic pancreatic cancer reveals site-specific immune regulatory pathways. Genome biology, 22(1):154, 2021.

Irizarry, R. A., Ladd-Acosta, C., Wen, B., Wu, Z., Montano, C., Onyango, P., Cui, H., Gabo, K., Rongione, M., Webster, M., et al. The human colon cancer methylome shows similar hypo-and hypermethylation at conserved tissue-specific cpg island shores. Nature Genetics, 41(2): 178–186, 2009.

Jeong, Y., Gerhäuser, C., Sauter, G., Schlomm, T., Rohr, K., and Lutsik, P. Methylbert enables read-level dna methylation pattern identification and tumour deconvolution using a transformer-based model. Nature Communications, 16(1):788, 2025.

Jolliffe, I. T. Principal component analysis for special types of data. Springer, 2002.

Jones, P. A. and Baylin, S. B. The epigenomics of cancer. Cell, 128(4):683–692, 2007.

Joshi, A., Boige, R., Zamparo, L., Tanielian, U., Garau-Luis, J. J., Chatzianastasis, M., Pandey, P., Sielemann, J., Seifert, A., Brand, M., et al. A long range foundation model for zero-shot predictions in single-cell and spatial transcriptomics data. 2025.

Katzman, J. L., Shaham, U., Cloninger, A., Bates, J., Jiang, T., and Kluger, Y. Deepsurv: personalized treatment recommender system using a cox proportional hazards deep neural network. BMC medical research methodology, 18: 1–12, 2018.

Kingma, D. P. and Ba, J. Adam: A method for stochastic optimization. arXiv preprint arXiv:1412.6980, 2014.

Kingma, D. P. and Welling, M. Auto-encoding variational bayes. arXiv preprint arXiv:1312.6114, 2013.

Klambauer, G., Unterthiner, T., Mayr, A., and Hochreiter, S. Self-normalizing neural networks. Advances in neural information processing systems, 30, 2017.

Krishna, G., Dharur, S., Rudovic, O., Dighe, P., Adya, S., Abdelaziz, A. H., and Tewfik, A. H. Modality drop-out for multimodal device directed speech detection using verbal and non-verbal features. In ICASSP 2024-2024 IEEE International Conference on Acoustics, Speech and Signal Processing (ICASSP), pp. 8240–8244. IEEE, 2024.

Lee, D. D. and Seung, H. S. Learning the parts of objects by non-negative matrix factorization. nature, 401(6755): 788–791, 1999.

Li, Z.-h., Hu, P.-h., Tu, J.-h., and Yu, N.-s. Luminal b breast cancer: patterns of recurrence and clinical outcome. Oncotarget, 7(40):65024, 2016.

Linder, J., Srivastava, D., Yuan, H., Agarwal, V., and Kelley, D. R. Predicting rna-seq coverage from dna sequence as a unifying model of gene regulation. Nature Genetics, pp. 1–13, 2025.

Liu, H., Tam, D., Muqeeth, M., Mohta, J., Huang, T., Bansal, M., and Raffel, C. A. Few-shot parameter-efficient finetuning is better and cheaper than in-context learning. Advances in Neural Information Processing Systems, 35: 1950–1965, 2022.

Lundberg, S. M. and Lee, S.-I. A unified approach to interpreting model predictions. Advances in neural information processing systems, 30, 2017.

Ma, M., Ren, J., Zhao, L., Tulyakov, S., Wu, C., and Peng, X. Smil: Multimodal learning with severely missing modality. In Proceedings of the AAAI Conference on Artificial Intelligence, volume 35, pp. 2302–2310, 2021.

Ma, M., Ren, J., Zhao, L., Testuggine, D., and Peng, X. Are multimodal transformers robust to missing modality? In Proceedings of the IEEE/CVF conference on computer vision and pattern recognition, pp. 18177–18186, 2022.

Ma, S., Zeng, A. G., Haibe-Kains, B., Goldenberg, A., Dick, J. E., and Wang, B. Integrate any omics: Towards genome-wide data integration for patient stratification. arXiv preprint arXiv:2401.07937, 2024.

Meng, X., Li, X., Yang, Q., Dai, H., Qiao, L., Ding, H., Hao, L., and Wang, X. Gene-moe: A sparsely gated prognosis and classification framework exploiting pan-cancer genomic information. arXiv preprint arXiv:2311.17401, 2023.

Nezakati, N., Reza, M. K., Patil, A., Solh, M., and Asif, M. S. Mmp: Towards robust multi-modal learning with masked modality projection. arXiv preprint arXiv:2410.03010, 2024.

Parker, J. S., Mullins, M., Cheang, M. C., Leung, S., Voduc, D., Vickery, T., Davies, S., Fauron, C., He, X., Hu, Z., et al. Supervised risk predictor of breast cancer based on intrinsic subtypes. Journal of clinical oncology, 27(8): 1160–1167, 2009.

Popov, M., Kallala, A., Ramesh, A., Hennouni, N., Khaitan, S., Gentry, R., and Cohen, A.-S. Leveraging state space models in long range genomics. arXiv preprint arXiv:2504.06304, 2025.

Ramazanova, M., Pardo, A., Ghanem, B., and Alfarra, M. Test-time adaptation for combating missing modalities in egocentric videos. In The Thirteenth International Conference on Learning Representations, 2025.

Saha, P., Mishra, D., Wagner, F., Kamnitsas, K., and Noble, J. A. Examining modality incongruity in multimodal federated learning for medical vision and language-based disease detection. arXiv preprint arXiv:2402.05294, 2024.

Sánchez, A., Fernández-Real, J., Vegas, E., Carmona, F., Amar, J., Burcelin, R., Serino, M., Tinahones, F., de Villa, M. C. R., Minãrro, A., et al. Multivariate methods for the integration and visualization of omics data. In Bioinformatics for Personalized Medicine: 10th Spanish Symposium, JBI 2010, Torremolinos, Spain, October 27-29, 2010. Revised Selected Papers, pp. 29–41. Springer, 2012.

Sandoval, J., Heyn, H., Moran, S., Serra-Musach, J., Pujana, M. A., Bibikova, M., and Esteller, M. Validation of a dna methylation microarray for 450,000 cpg sites in the human genome. Epigenetics, 6(6):692–702, 2011.

Scarselli, F., Gori, M., Tsoi, A. C., Hagenbuchner, M., and Monfardini, G. The graph neural network model. IEEE transactions on neural networks, 20(1):61–80, 2008.

Shannon, C. E. A mathematical theory of communication. The Bell system technical journal, 27(3):379–423, 1948.

Srivastava, N., Hinton, G., Krizhevsky, A., Sutskever, I., and Salakhutdinov, R. Dropout: a simple way to prevent neural networks from overfitting. The journal of machine learning research, 15(1):1929–1958, 2014.

Stark, R., Grzelak, M., and Hadfield, J. Rna sequencing: the teenage years. Nature Reviews Genetics, 20(11):631–656, 2019.

Stouffer, S. A., Suchman, E. A., DeVinney, L. C., Star, S. A., and Williams Jr, R. M. The american soldier: Adjustment during army life.(studies in social psychology in world war ii), vol. 1. 1949.

Subramanian, A., Tamayo, P., Mootha, V. K., Mukherjee, S., Ebert, B. L., Gillette, M. A., Paulovich, A., Pomeroy, S. L., Golub, T. R., Lander, E. S., et al. Gene set enrichment analysis: a knowledge-based approach for interpreting genome-wide expression profiles. Proceedings of the National Academy of Sciences, 102(43):15545–15550, 2005.

Tanvir, R. B., Islam, M. M., Sobhan, M., Luo, D., and Mondal, A. M. Mogat: a multi-omics integration framework using graph attention networks for cancer subtype prediction. International Journal of Molecular Sciences, 25(5): 2788, 2024.

Traag, V. A., Waltman, L., and Van Eck, N. J. From louvain to leiden: guaranteeing well-connected communities. Scientific reports, 9(1):1–12, 2019.

Tran, B. and Bedard, P. L. Luminal-b breast cancer and novel therapeutic targets. Breast Cancer Research, 13(6): 221, 2011.

Vale-Silva, L. A. and Rohr, K. Long-term cancer survival prediction using multimodal deep learning. Scientific Reports, 11(1):13505, 2021.

Vaswani, A., Shazeer, N., Parmar, N., Uszkoreit, J., Jones, L., Gomez, A. N., Kaiser, Ł., and Polosukhin, I. Attention is all you need. Advances in neural information processing systems, 30, 2017.

Verhaak, R. G., Tamayo, P., Yang, J.-Y., Hubbard, D., Zhang, H., Creighton, C. J., Fereday, S., Lawrence, M., Carter, S. L., Mermel, C. H., et al. Prognostically relevant gene signatures of high-grade serous ovarian carcinoma. The Journal of clinical investigation, 123(1), 2012.

Wagner, J., Andre, E., Lingenfelser, F., and Kim, J. Exploring fusion methods for multimodal emotion recognition with missing data. IEEE Transactions on Affective Computing, 2(4):206–218, 2011.

Waqas, A., Tripathi, A., Ahmed, S., Mukund, A., Farooq, H., Schabath, M. B., Stewart, P., Naeini, M., and Rasool, G. Self-normalizing foundation model for enhanced multi-omics data analysis in oncology. arXiv preprint arXiv:2405.08226, 2024.

Weber, M., Hellmann, I., Stadler, M. B., Ramos, L., Pääbo, S., Rebhan, M., and Schübeler, D. Distribution, silencing potential and evolutionary impact of promoter dna methylation in the human genome. Nature Genetics, 39 (4):457–466, 2007.

Yang, F., Wang, W., Wang, F., Fang, Y., Tang, D., Huang, J., Lu, H., and Yao, J. scbert as a large-scale pretrained deep language model for cell type annotation of single-cell rnaseq data. Nature Machine Intelligence, 4(10):852–866, 2022.

Zhang, W., Xu, D., Zhang, J., and Ouyang, W. Progressive modality cooperation for multi-modality domain adaptation. IEEE Transactions on Image Processing, 30:3293–3306, 2021a.

Zhang, X., Xing, Y., Sun, K., and Guo, Y. Omiembed: a unified multi-task deep learning framework for multiomics data. Cancers, 13(12):3047, 2021b.

Zhi, Z., Liu, Z., Elbadawi, M., Daneshmend, A., Orlu, M., Basit, A., Demosthenous, A., and Rodrigues, M. Borrowing treasures from neighbors: In-context learning for multimodal learning with missing modalities and data scarcity. arXiv preprint arXiv:2403.09428, 2024.

